# Exponential Growth of Glioblastoma *In Vivo* Driven by Rapidly Dividing and Outwardly Migrating Cancer Stem Cells

**DOI:** 10.1101/723601

**Authors:** Lisa Buchauer, Muhammad Amir Khan, Yue Zhuo, Chunxuan Shao, Peng Zou, Weijun Feng, Mengran Qian, Gözde Bekki, Charlotte Bunne, Anna Neuerburg, Azer Aylin Acikgöz, Mona Tomaschko, Zhe Zhu, Heike Alter, Katharina Hartmann, Olga Friesen, Klaus Hexel, Thomas Höfer, Hai-Kun Liu

## Abstract

How overall tumor growth emerges from the properties of functionally heterogeneous tumor cell subpopulations is a fundamental question of cancer biology. Here we combined lineage tracing, continuous monitoring of tumor mass, proliferation assays and transcriptomics with mathematical modeling and statistical inference to dissect the growth of glioblastoma in mice. We found that tumors grow exponentially at the rate of symmetric divisions of brain tumor stem cells (BTSCs). Spatial modeling predicts, and data show, that BTSCs accumulate at the tumor rim rather than in the core. The physiological differentiation hierarchy downstream of BTSCs is preserved in mice and humans: transit amplifying progenitors give rise to terminally differentiated cells. Consistent with our quantification of the mechanisms underlying tumor growth, molecular data show elevated expression of cell cycle-and migration-related genes in BTSCs. Our systematic approach reveals fundamental properties of glioblastoma and may be transferable to the study of other animal models of cancer.

## Introduction

The growth laws of human tumors have been intensively studied since the early days of cancer research, and it was hoped that insight into tumor growth dynamics would help optimize treatment schedules (see Collins et al. 1956 for an influential early paper and Byrne, 2010 for comprehensive review) [1,2]. Thus far, however, quantitative studies of tumor growth have remained largely phenomenological, and the integration from single cell fate decisions to tumor growth remains challenging. Recent work has begun to connect clonal genetic evolution and tumor growth [3–5]. However, a complementary quantitative view on the proliferation, survival and differentiation properties of single tumor cells *in vivo*, and how these impact overall tumor expansion, is still missing.

A key insight from the large body of recent molecular and mechanistic work is that tumors are intrinsically heterogeneous at the single-cell level, with respect to both oncogenic driver mutations as well as cell behavior in terms of proliferation and survival in response to treatment [6–11]. The cancer stem cell (CSC) hypothesis suggests that functional tumor cell heterogeneity arises, in large part, from perturbations of the normal stem and progenitor cell hierarchy seen in renewal tissues [7,12–14]. However, it has been difficult to study the growth dynamics of such hierarchical tumors in vivo. Intratumoral genetic heterogeneity may be dissected by genome sequencing, yet this provides no direct information on variability in cell behavior and, moreover, may be hampered by limited sampling of human tumors [9]. Recent studies using lineage tracing in animal models provide more direct evidence of existence and significance of CSC-like cells in mouse tumors and human xenograft tumors [15–19]. However, we lack quantitative understanding of tumor growth and treatment response in hierarchically organized tumors arising from CSCs. In particular, we do not know how the stem-cell and non-stem cell fractions of the tumor contribute to its overall growth and how they are spatially organized within the tumor.

To address these questions, we here focus on a mouse model of a fast-growing tumor, glioblastoma multiforme (GBM). GBM is one of the most aggressive cancers and is associated with poor prognosis [20]. To date, there is no effective therapy despite rapid development in molecular classification of GBM [21]. Heterogeneity at the genetic, cell-phenotypic and morphological levels both within individual tumors and across different tumors is a hallmark of GBM. The existence of brain tumor stem cells (BTSCs) [22–24], which are reported to be resistant to conventional therapy [16–25], is tightly linked to this property. We have previously shown that the nuclear receptor Tlx (Nr2e1) is essential for neural stem cell (NSC) maintenance and tumor initiation [26–28]. Recently, we demonstrated that Tlx^+^ cells behave as stem cells in primary mouse GBM, and that removal of Tlx in BTSCs leads to prolonged survival of tumor-bearing mice [18]. Here, we combine mouse genetic tools with mathematical modeling and statistical inference to define how individual tumor stem cells and their progeny shape bulk growth and treatment response of GBM *in vivo*.

## Results

### Exponential Growth of Brain Tumors Is Enabled by Cancer Stem Cell Migration

To follow tumor growth in unperturbed primary GBM, we used a mouse model based on the RCAS/Nestin-tv-a (Ntv-a) transgenic system [29], overexpressing Pdgfb (platelet-derived growth factor subunit B) and Akt to efficiently induce high-grade gliomas [18]. We also delivered RCAS-Firefly Luciferase (Fluc) at tumor induction along with Pdgfb and Akt to quantify tumor cell number over time (Figure 1A). We imaged 23 tumor-bearing animals using this approach and fit the most widely used growth laws for solid tumor growth – exponential growth, Gompertzian growth and linear radial growth [30–33] – to the data (*Supplementary Theory* Figures T1-T3). Initially tumors may grow exponentially before growth slows due to limitations in space or supply of nutrients or oxygen; both Gompertzian and linear radial growth laws implement growth limitation with increasing tumor size. Model selection based on Akaike’s information criterion, which includes a penalty term that rises with the number of model parameters [34], showed that for the majority of tumors (18/23, or 78%), an exponential increase of cell numbers over the imaging period described the data best, while the remainder was fit best by Gompertzian growth (*Supplementary Theory* Figure T1-T3), suggesting that the majority of tumors grew unhindered by spatial or nutrient constraints. The exponential growth rates of individual tumors varied over a 4.6-fold range with an average growth rate of (0.21 ± 0.02) / day (mean ± SEM), equivalent to an average doubling time of 3.3 days (Figure 1B).

**Figure 1.**
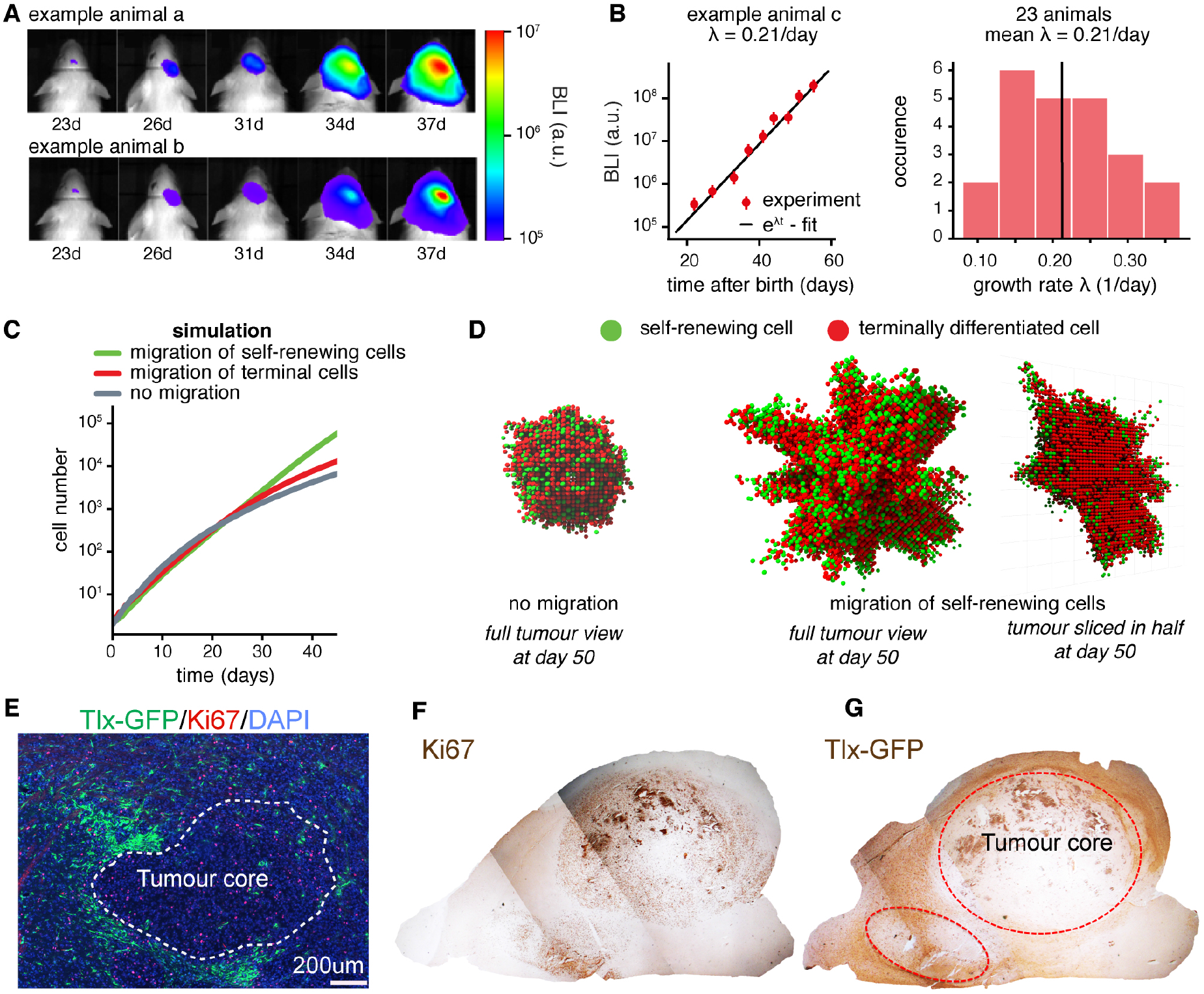
BTSCs accumulate in the shell of exponentially growing brain tumors. **A**, Bioluminescence imaging of tumor growth in two exemplary N-tva mice. **B**, Left: Bioluminescence imaging (BLI) of tumor growth in one exemplary N-tva mouse and exponential growth-law fit yielding a growth rate of 0.21/day. Right: Histogram of growth rates derived from exponential fits to 23 mice, yielding a mean growth rate of 0.21/day. For all 23 fit results, see *Supplementary Theory*. **C**, Modelled tumor growth curves using a 3D-grid based simulation of self-renewing cells and terminally differentiated cells and one of three migration scenarios. Curves show mean of 50 stochastic simulations. For simulation algorithm and parameters see *Supplementary Theory*. **D**, Simulated example tumors at day 50 after initiation. Left: Tumor developed without cellular migration. Middle and right: Tumor developed with outwards migration of self-renewing cells. The right panel shows a cut through the tumor shown in the middle panel: self-renewing cells accumulate in the tumor shell. **E**, Tlx-GFP/Ki67 staining of early tumors shows a tumor border distribution of Tlx-GFP cells. **F**&**G**, Ki67(F) and Tlx-GFP(G) staining of Tlx-GFP xenograft tumor show that Tlx-GFP^+^ cells are distributed around the tumor border.

To interrogate how the observed exponential growth of GBM may occur, we developed a simple grid-based stochastic simulation, with self-renewing tumor cells (BTSCs) dividing into either two BTSCs or one BTSC and one differentiated, non-proliferative progenitor cell. For cells to divide there must be enough space in their surroundings to accommodate the two daughter cells (see *Supplementary Theory* and simulation code for full details). When cells stayed close to their place of birth, the growth rate decreased as the tumor became larger (Figure 1C, gray curve), resulting in compact, near-spherical tumors (Figure 1D, left panel). When non-proliferative progeny was allowed to migrate, the slow-down of tumor growth was attenuated but not prevented (Figure 1C, red line). By contrast, exponential tumor growth was sustained when we allowed BTSCs to migrate away from the center of the tumor mass (Figure 1C, green line). Migration of BTSCs caused a fuzzy appearance of the tumor with invading fingers (Figure 1D, middle panel). Cross-sections through these simulated tumors showed that migratory BTSCs are preferentially distributed around the tumor border (Figure 1D, right panel). Thus, our simulation of GBM growth suggests that exponential growth can be sustained even for large tumors when self-renewing cells disperse outwards and hence become localized to the tumor margin.

To test this prediction, we analyzed brain tumors from Tlx-GFP;N-tva mice, for which we showed previously that Tlx-GFP marks BTSCs [18]. We found that Tlx-GFP positive cells were preferentially distributed around the tumor margin (Figure 1E). To exclude potential contamination by Tlx-GFP-expressing resident NSCs, we orthotopically transplanted the Tlx-GFP BTSCs into NOD/SCID mice and found a similar Tlx-GFP distribution pattern (Figure 1F,G). Both simulation and experiment suggest that cells with proliferative potential migrate towards the tumor edges and invade surrounding brain tissue. This finding may provide a rationale for the observation that most recurrences of human glioblastomas occur at the resection margin [35].

### Lineage Tracing Reveals a BTSC-Based Differentiation Hierarchy in Tumors

To probe the cellular hierarchy within the tumor at the level of individual BTSC clones, we next performed lineage tracing of Tlx-positive BTSCs using Tlx-CreER^T2^;Ntv-a;Confetti mice [35]. Upon tamoxifen (TAM) mediated Cre activation, this mouse model allows the random labeling of BTSCs with one of four colors (blue, green, red or yellow) [15]. We induced sparse confetti color labeling of single Tlx-GFP cells in Tlx-GFP;Tlx-CreER^T2^;Ntv-a;Confetti mice (Figure 2A). Focusing on the red (RFP) clones, we ascertained that the initial tracing started with Tlx-GFP-positive BTSCs (Figure 2B). We observed that clones contained both Tlx-GFP positive and negative cells, and 20 days after TAM application, the bulk of the RFP population consisted of Tlx-GFP-negative cells (Figure 2C). Hence Tlx-expressing BTSCs give rise to non-Tlx expressing progeny. Tlx-GFP-positive cells were negative for Olig2 (Figure 2D), which was suggested to be a BTSC marker [23]. However, the progenies of Tlx-expressing cells are positive for Olig2 (Figure 2E), suggesting that Tlx-expressing cells give rise to Tlx^-^/Olig2^+^ cells in vivo, and placing Olig2 expression downstream of the top BTSCs, as a marker of tumor progenitor cells. These results indicate the persistence of a differentiation hierarchy in high-grade brain tumors.

**Figure 2.**
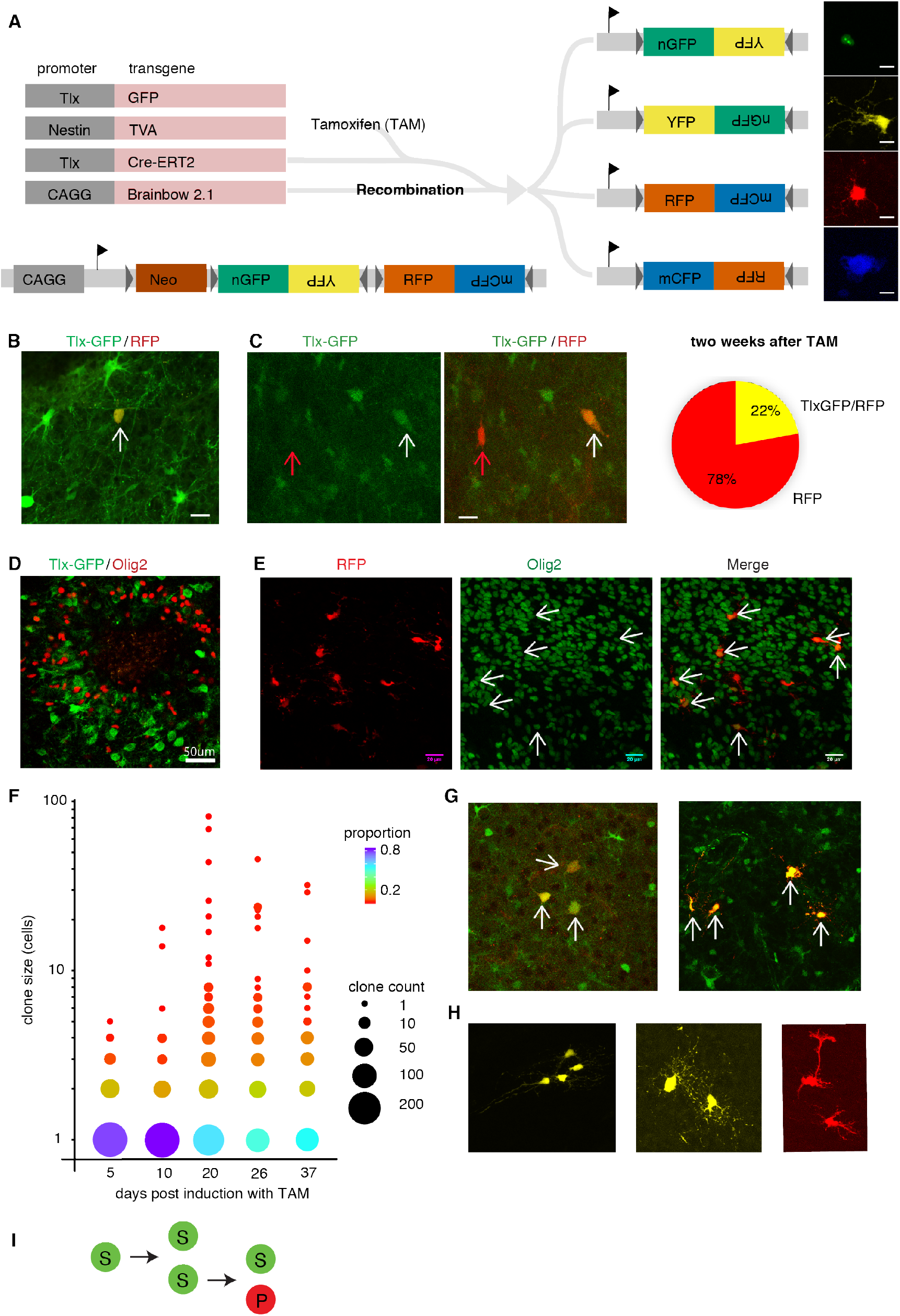
Single cell tracing of BTSCs reveals cellular differentiation hierarchy with morphologically heterogeneous progenitors. **A**, Schematic illustration of clonal tracing of Tlx-positive tumor cells in Tlx-CreER^T2^;Ntv-a;Confetti mice. **B**, Singlecell induction of confetti labeling in Tlx-GFP^+^ cells. The arrow indicates a single Tlx-GFP^+^ cell labeled with RFP after TAM injection. Scale bar: 20μm if not indicated otherwise. **C**, Asymmetric division was observed during tracing; arrows indicate a Tlx-GFP^+^/RFP^+^ cell (white arrow) giving rise to a Tlx-GFP^−^cell (red arrow). The pie chart shows the proportions of Tlx-GFP^+^ and Tlx-GFP^-^cells 20 days after TAM injection within all analyzed red clones. **D**, Tlx-GFP^+^ cells are negative for Olig2 in mouse brain tumors. **E**, An RFP^+^ clone derived from a Tlx^+^ cell stained for RFP and Olig2; arrows indicate double positive cells. **F**, Summary of clonal tracing results over time. While the fraction of large clones increases during the first three weeks, a high number of singlecell clones was found at all time points. **G**, Examples for clones containing several Tlx-GFP^+^ positive cells only, suggesting that symmetric division occurred to expand the Tlx-GFP^+^ population *in vivo*. **H**, Labelled tumour cells and their progenitors display morphological heterogeneity. **I**, Summary of the above for model construction. *S* – brain tumor stem cell, *P* – tumor progenitor cell.

We then performed a series of clonal tracing experiments starting from single Tlx-expressing tumor cells in Tlx-CreER^T2^;Ntv-a;Confetti tumor-bearing mice. The mean size of traced clones increased in the first three weeks and then plateaued, with individual clone size being heterogeneous (Figure 2F). Intriguingly, throughout local areas, we observed small clones, including those made up of only a single cell (Figure 2E). Two effects may explain this phenomenon. On the one hand, a fraction of BTSCs could be dormant. On the other hand, BTSCs may migrate out of the analyzed area and found new clones at a distance. Three-dimensional reconstruction of clones across multiple microscopy sections showed local but partially overlapping clones, confirming that BTSCs are migratory (Figure S1). We found that some multicellular clones only consisted of Tlx-GFP^+^ cells (Figure 2G), suggesting that symmetric expansion of BTSCs occurred in such clones. In other clones, the tumor cells derived from Tlx^+^ BTSCs showed varied morphologies characteristic of cell differentiation (Figure 2H). To develop a cell-based quantitative model of tumor growth, we conclude that symmetric BTSC divisions expanding BTSCs and asymmetric divisions giving rise to more differentiated progeny co-occur in individual clones (Figure 2I).

### Tumor Progenitor Cells Do Not Self-Renew

Our clonal tracing data suggest that BTSCs differentiate into tumor progenitor cells (TPCs). However, non-stem cells might secondarily acquire CSC properties [12]. To test for this, we traced the fate of the proliferative Tlx-GFP-negative population by injecting the tumor-bearing Tlx-GFP;Ntv-a mice with a retrovirus carrying a dsRed-expressing construct, which infected Tlx-GFP^−^ cells downstream of BTSCs. Most of the dsRed-labelled cells expressed Ki67 one day after labeling (Figure 3A), consistent with retroviral infection occurring in rapidly dividing cells. However, these cells lost Ki67 expression over time, with the fraction of Ki67^+^ dropping to about 10% 13 days after labelling (Figure 3B). These results suggest that the fast-dividing Tlx-GFP^−^ cells do not sustain their proliferative capacity. The dsRed-labeled population is positive for Olig2 (Figure 3C), again indicating Olig2 as a marker for intermediate tumor progenitors in vivo. Importantly, we did not observe any dsRed^+^ cells expressing Tlx-GFP during the entire period of analysis (Figure 3D). This implies that, first, BTSCs are initially not targeted by the retrovirus and, second, dedifferentiation of tumor progenitor cells into BTSCs did not occur (or at best occured so rarely as not to be picked up in our data) during tumor progression. We also observed morphological heterogeneity within the dsRed^+^ cells, characteristic of differentiated cell fates (Figure 3E). Moreover, tumor cells formed organized structures by extending cell protrusions towards the same niche, which may indicate (chemotactically) regulated migration (Figure 3F). Taken together, these data show that TPCs do not revert to a BTSC phenotype during GBM growth but, rather, have a limited proliferative capacity and eventually give rise to non-proliferative differentiated tumor cells (DTCs, Figure 3G).

**Figure 3.**
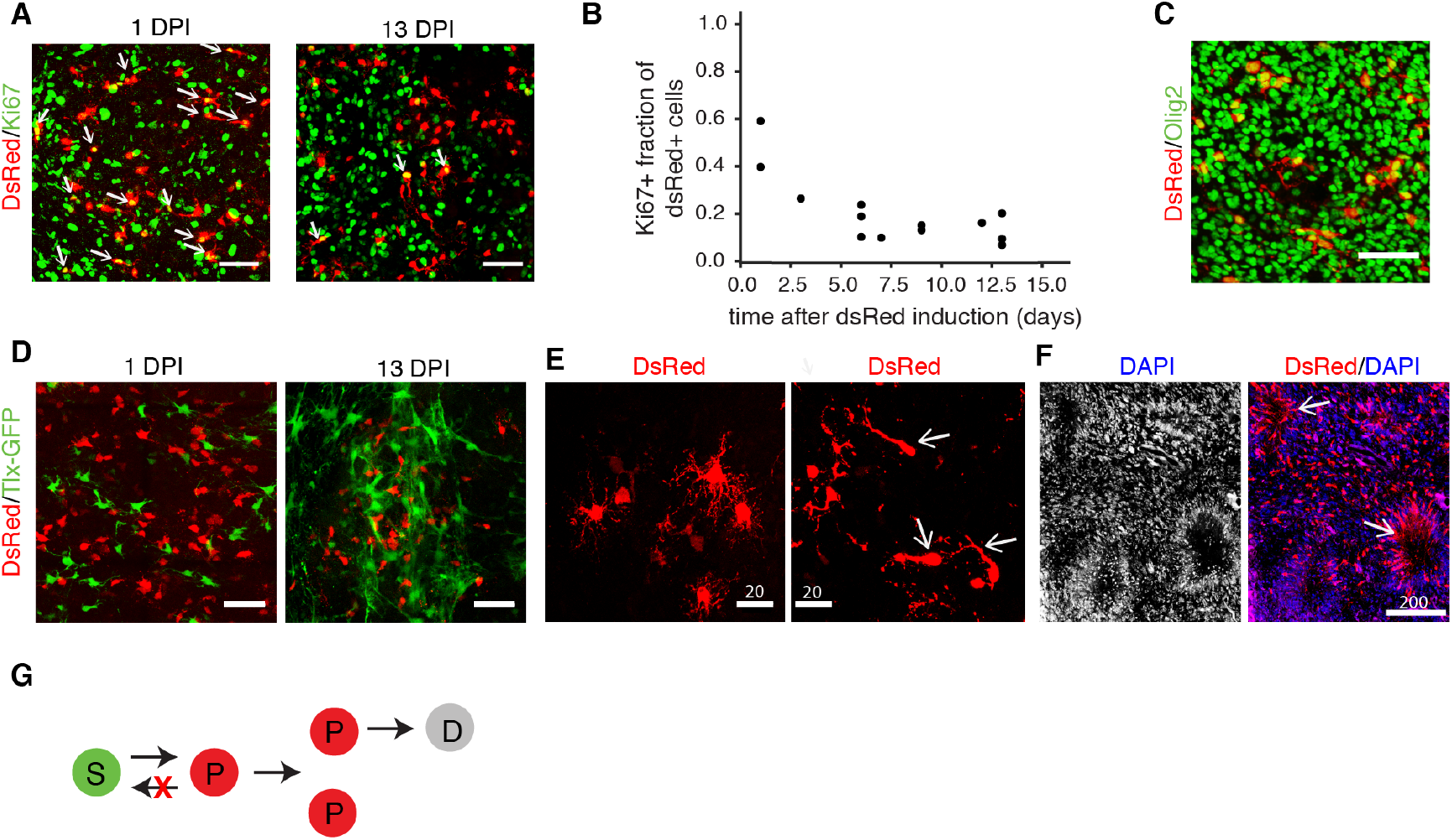
Retroviral tracing of rapidly dividing tumor progenitor cells supports their intermediate position in the differentiation hierarchy. **A**, Retroviral labeling of tumor cells with dsRed. dsRed-labeled cells are mostly positive for Ki67 one day after label induction and mostly negative for Ki67 13 days after induction; arrows indicate double positive cells. **B**, Quantification shows a continuous decline in the proliferation index of dsRed-labeled cells over time. Dots represent individual mice. **C**, dsRed-labeled cells are positive for Olig2. **D**, Retrovirally labeled dsRed cells in Tlx-GFP tumor mice are negative for Tlx-GFP 1 day after induction (DPI), and they do not gain expression of Tlx-GFP over time (13DPI). **E** & **F**, Morphological heterogeneity of between dsRed-labelled tumor cells. **G**, Summary of the above for model construction. *S* – brain tumor stem cell, *P* – tumor progenitor cell, *D* - differentiated tumor cell.

### Mosaic Analysis Shows Continuous Cellular Turnover

The dsRed-tracing results suggest that GBM growth *in vivo* is characterized by cellular turnover during which initially dividing cells lose their proliferation capacity and are replaced by new actively dividing cells. To analyze the dynamics of the different cell populations involved in this process, we adopted a mosaic labeling approach based on the FLEx system [36] to simultaneously trace clones derived from BTSCs and TPCs (Figure 4A). The RGFlex construct was delivered via the RCAS vector during tumor induction, resulting in an initial RFP labeling of tumor cells. Tlx-CreER^T2^;Ntv-a mice were used for this experiment to allow, upon TAM injection, simultaneous labeling of stem-cell-driven expansion with GFP and non-stem-cell-driven expansion with RFP (Figure 4A). TAM was given on ten consecutive days to achieve complete labeling of all Tlx-CreER^T2^ cells in the mouse tumor; times reported here are to be read as following the end of the 10-day TAM administration phase. Two days after TAM application, we observed GFP expression in brain tumors. The GFP-expressing cells were found preferentially around the tumor border, with moderate levels of Ki67 expression (Figure 4B). We detected RFP signal in some GFP^+^ cells, indicating a recent conversion from RFP to GFP. We also observed migratory GFP^+^ cells (Figure 4C). These data confirm that Tlx^+^ BTSCs accumulate at the tumor margin. Of note, GFP^+^ cells around the tumor were highly polarized and did not form clusters (Figure 4D), which is unlike collective neuroblast migration in the adult SVZ olfactory bulb system [37,38].

**Figure 4.**
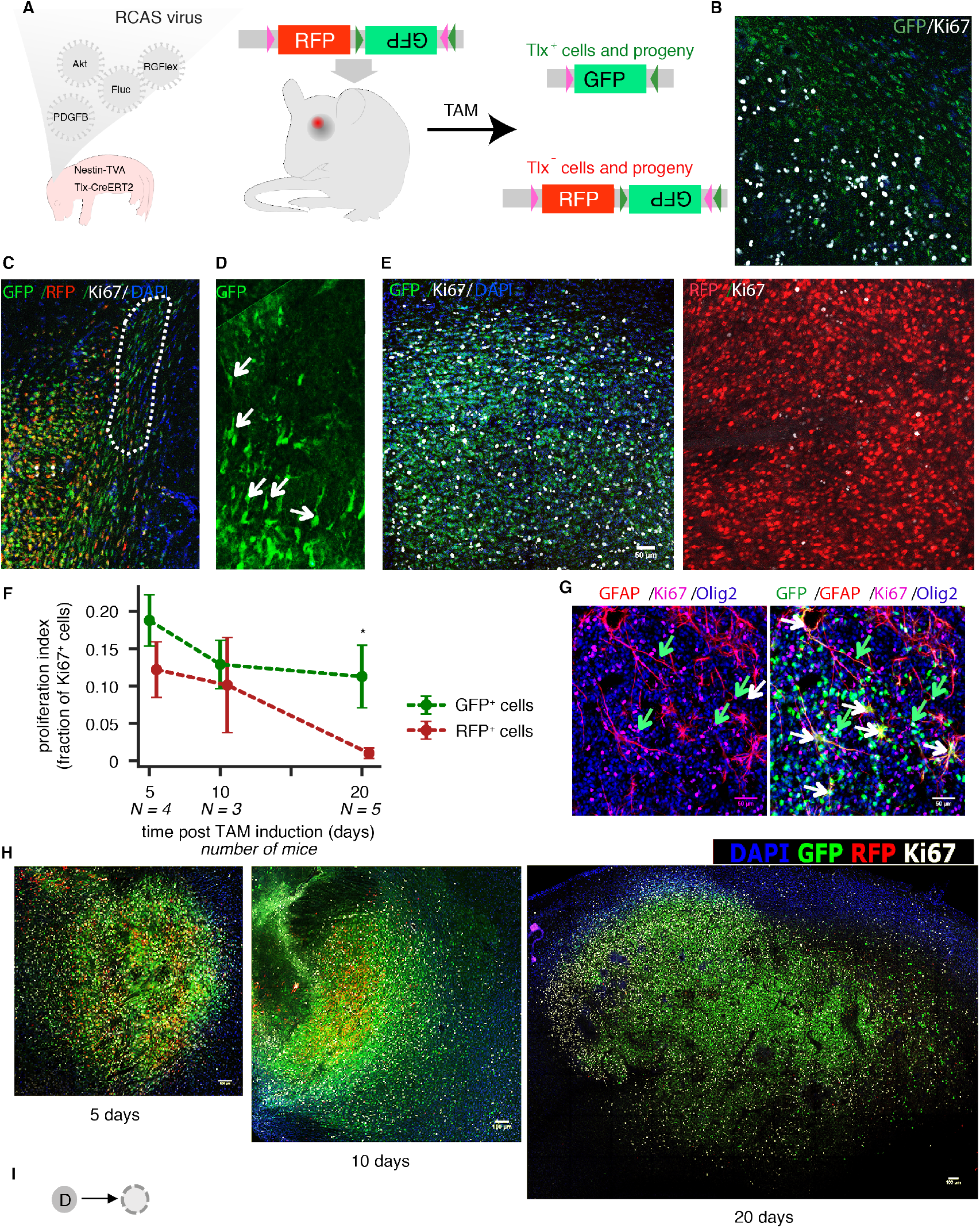
Mosaic analysis of BTSCs and non-BTSCs allows visualization of cellular turnover during tumor progression. **A**, Experimental design of a Flex-based tracing strategy. After TAM induction of Tlx-CreERT2 mediated color conversion, BTSCs and their progeny are labeled with GFP; the non-BTSCs and their progeny remain labeled with RFP. **B**, GFP labeled cells are found at the tumor border 2 days after TAM treatment. Ki67 (white) indicates tumor bulk. **C**, A GFP/RFP double-positive area indicates that color conversion has occurred recently. GFP^+^ cells are found in the corpus callosum with a migratory pattern (marked area). **D**, GFP^+^ cells show unipolar morphology (arrows), non-clustered but with shared orientation, suggesting chemotaxis-mediated cellular migration. **E**, Ki67 staining shows that GFP positive clones are still proliferative 20 days after TAM, whereas the RFP clones are less proliferative. **F**, Quantitative analysis of proliferation index suggests RFP clones do not retain proliferation potential. Number of mice evaluated per time point is indicated, means are plotted. Error bars show standard error of mean. Horizontal offset between RFP and GFP points is for visibility only. Proliferation index at day 20 differs significantly between the groups (double-sided t-test, p=0.04). **G**, GFP co-staining with GFAP and Olig2 suggests that Tlx-positive cells generate both astrocytes (white arrows) and oligodendrocytes (green arrows). **H**, GFP/RFP/Ki67 staining shows a gradual increase of the GFP^+^ tumor fraction. **I**, Summary of the above for model construction – DTCs (*D*) disappear from the tumor.

While GFP^+^ cells showed moderate proliferation as judged by Ki67 staining two days after color conversion (Figure 4B), high levels of Ki67 expression were observed 20 days after conversion suggesting a high average proliferation rate of active BTSCs and their immediate progenitors (Figure 4E). In agreement with the retrovirus-mediated tracing experiment, RFP^+^ cells were less proliferative than GFP^+^ cells 20 days after color conversion (Figures 4E, F). The GFP^+^ population was found to contain both astrocyte-and oligodendrocyte-like tumor cells 20 days after color conversion, confirming the existence of functional heterogeneity between tumor cells (Figure 4G).

Of note, the fraction of RFP^+^ cells within all labelled cells was decreasing as a function of time after color conversion with the tumor becoming increasingly occupied by GFP^+^ cells (Figure 4H). This suggests that not only do TPCs lose their proliferation potential while differentiating into DTCs, but DTCs are lost from the tumor via cellular turnover within a relatively short time frame during a phase in which the overall tumor mass is growing exponentially. To incorporate this finite lifetime of progenitor cells into the tumor growth model, we added a cell death process of non-proliferating progenitors whereby they are eventually lost from the system (Figure 4I).

### Mathematical Model of the Cellular Hierarchy and Brain Tumor Growth

Step by step (Figure 2–4), we have elucidated a hierarchical model of brain tumor growth consisting of migratory and proliferative brain tumor stem cells (BTSCs, model: *S*), proliferating tumor progenitor cells (TPCs, model: *P*), and exhausted progenitors or differentiated tumor cells (DTCs, model: *D*) (Figure 5A). To integrate the experimental data sets into a quantitative model of brain tumor growth, we translated the scheme into a system of ordinary differential equations (Figure 5B). We assumed, without loss of generality, that the differentiating events from BTSCs to TPCs occur by asymmetric divisions. Keeping the model as simple as possible to facilitate parameter inference, we did not explicitly describe cell migration in space but assumed that BTSCs migrate to outward locations where they can continue to divide.

**Figure 5.**
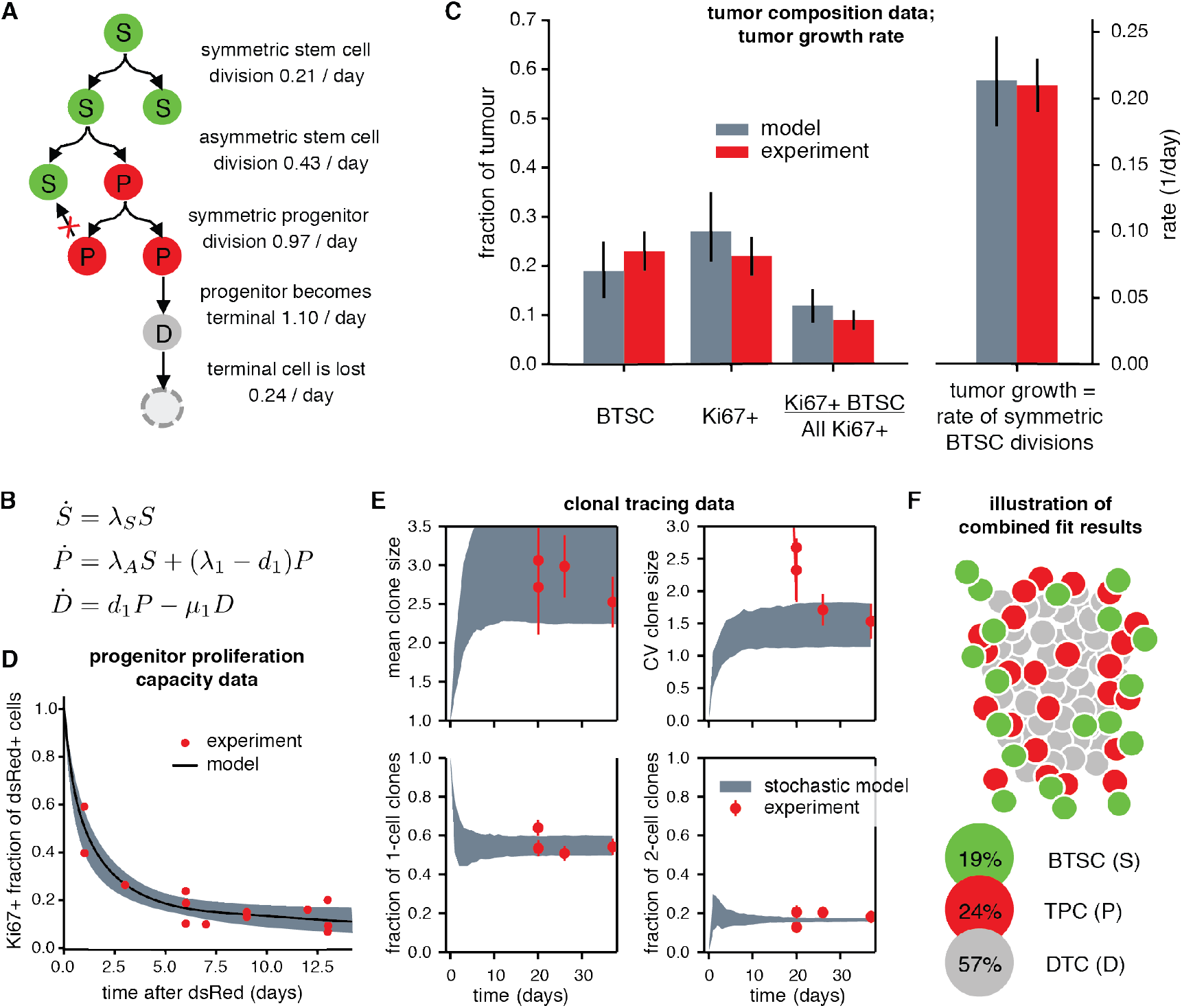
Estimation of cellular rates and tumor composition using a hierarchical mathematical model of glioblastoma growth. **A**, Hierarchical tumor growth model built from qualitative experimental observations as given in Figures 2I, 3G and 4I with *maximum a posteriori* probability parameters estimated by simultaneous incorporation of experimental data in panels C and D. **B**, Translation of the model shown in panel **A** into a set of coupled ordinary differential equations. **C** & **D**, Experimental evidence used for parameter estimation; comparison between experiment and model values. Model error bars in C and shaded-area in D show a=0.05 posterior probability bounds, see *Supplementary Theory*. Experimental error bars, where applicable, show standard error of mean. **E**, Fit of the model given in panel B complemented by a stochastic migration process (see *Supplementary Theory*) to statistics derived from single-cell clonal tracing (Figure 2). Shaded areas derived from 1000 independent model simulations with parameter sets within a=0.05 posterior probability bounds. Experimental errors derived by bootstrapping. CV - coefficient of variation. **F**, Sketch of tumor composition and architecture after incorporation of all experimental data. More than half of the tumor consists of DTCs; proliferative action (BTSCs and TPCs) is concentrated in the tumor shell.

A key consequence of this hierarchical cellular organization of the tumor is that the exponential growth rate of the tumor mass is equal to the symmetric division rate of the BTSCs (*Supplementary Theory*) [39]; the proliferation, differentiation and death parameters of TPCs and DTCs have no impact on the long-term growth of the tumor mass because they lack the ability to self-renew. Moreover, we find that during the exponential growth phase the relative proportions of BTSCs, TPCs and DTCs remain constant; these fractions are governed by all proliferation and death rates of the various subpopulations (*Supplementary Theory*). Hence the growing brain tumor has properties that depend only on the BTSCs (overall tumor growth rate) and properties that depend on all subpopulations (heterogeneous composition of the tumor).

Using a Bayesian parameter estimation framework (*Supplementary Theory*), we fitted the model simultaneously to the bioluminescence growth curves (Figure 1B), the fraction of proliferating progenitors as a function of time (Figure 3B), the fraction of Tlx-GFP-positive cells (Figure 2C) as well as previously published proliferation data [18,40]. In addition, we determined the proportion of BTSCs by FACS analysis of the Tlx-GFP population in tumors induced with PDFGB, Akt and RFP to be ~20% of the total tumor cells (Figures S2A & S2B). The model accounted for all these bulk experiments simultaneously (Figure C and D). In order to incorporate the data of the singlecell tracing experiment (Figure 2) into the model, we added a stochastic simulation module describing the tumor at the clonal level and estimated all parameters jointly using approximate Bayesian computation (*Supplementary Theory*). When BTSC migration was accounted for in this way, the model also explained the experimental distribution of single BTSC-derived tumor cell clones (Figure 2F), as summarized by its mean, variance and fraction of 1-and 2-cell clones (Figure 5E).

Overall, the data determined a unique parameter set (Figure 5A, *Supplementary Theory* Figures T4-T7), showing that BTSCs divide symmetrically once in five days while they produce TPCs via asymmetric division twice as often. Thus, BTSCs divide regularly and frequently during tumor progression. Note that these are average values for the BTSC population; if a fraction of the BTSCs is quiescent (e.g., during migration), the proliferative remainder will divide more frequently. Thus, at least a fraction of BTSCs is proliferatively very active. The doubling time of the BTSC population by symmetric division is ln 2 · 5 days = 3.5 days, which quantitatively matches the average doubling time of the tumor mass of 3.3 days observed by bioluminescence.

Lower down in the hierarchy, TPCs divide once per day and turn into DTCs only slightly faster. Consequently, DTCs make up more than half of the tumor mass while BTSCs occupy around 20% of the tumor mass (Figure 5F). TPC proliferation agrees with division rates of normal neural progenitors from the SVZ [41]. Due to the rapid differentiation of TPCs into DTCs and subsequent cell death, TPCs cells are unable to sustain tumor growth. Taken together, our data imply that TPCs display nearly normal proliferative behavior whereas BTSCs divide abnormally often when compared to normal neural stem cells of the SVZ and drive overall tumor growth.

### Targeting Brain Tumor Stem Cells Causes Tumor Regression

Standard chemotherapy with temozolomide (TMZ) has been designed to target proliferating tumor cells regardless of their place in the BTSC differentiation hierarchy. We simulated TMZ treatment by letting cell divisions during treatment cause cell death but otherwise keeping all parameters as estimated for tumor growth (Figs. 6A, B). For a treatment period of ten days, the model predicted a mean drop in tumor volume by factor ~50. This minimal tumor size is reached about five days after the end of the treatment, as the effect of killing BTSCs takes time to propagate through to TPCs and DTCs. However, not all BTSCs are eliminated and the tumor regrows driven by surviving BTSCs and reaches pre-treatment volume around 20 days after the end of the treatment (Figure 6C, *Supplementary Theory* Figure T7). To experimentally monitor the tumor response to TMZ treatment, we used the N-tva mice injected with Pdgfb, Akt and Fluc constructs. Once tumor size exceeded a threshold (1-5×10^7^ photons/sec), the mice were subjected to treatment. As predicted by the model, TMZ was very efficient in transiently reducing the tumor volume, with the minimum size reached with a similar delay as in the model after the end of treatment (Figure 6D, Figure S3A). Invariably, however, the tumors then regrew rapidly, congruent with model predictions (Figure 6D).

**Figure 6.**
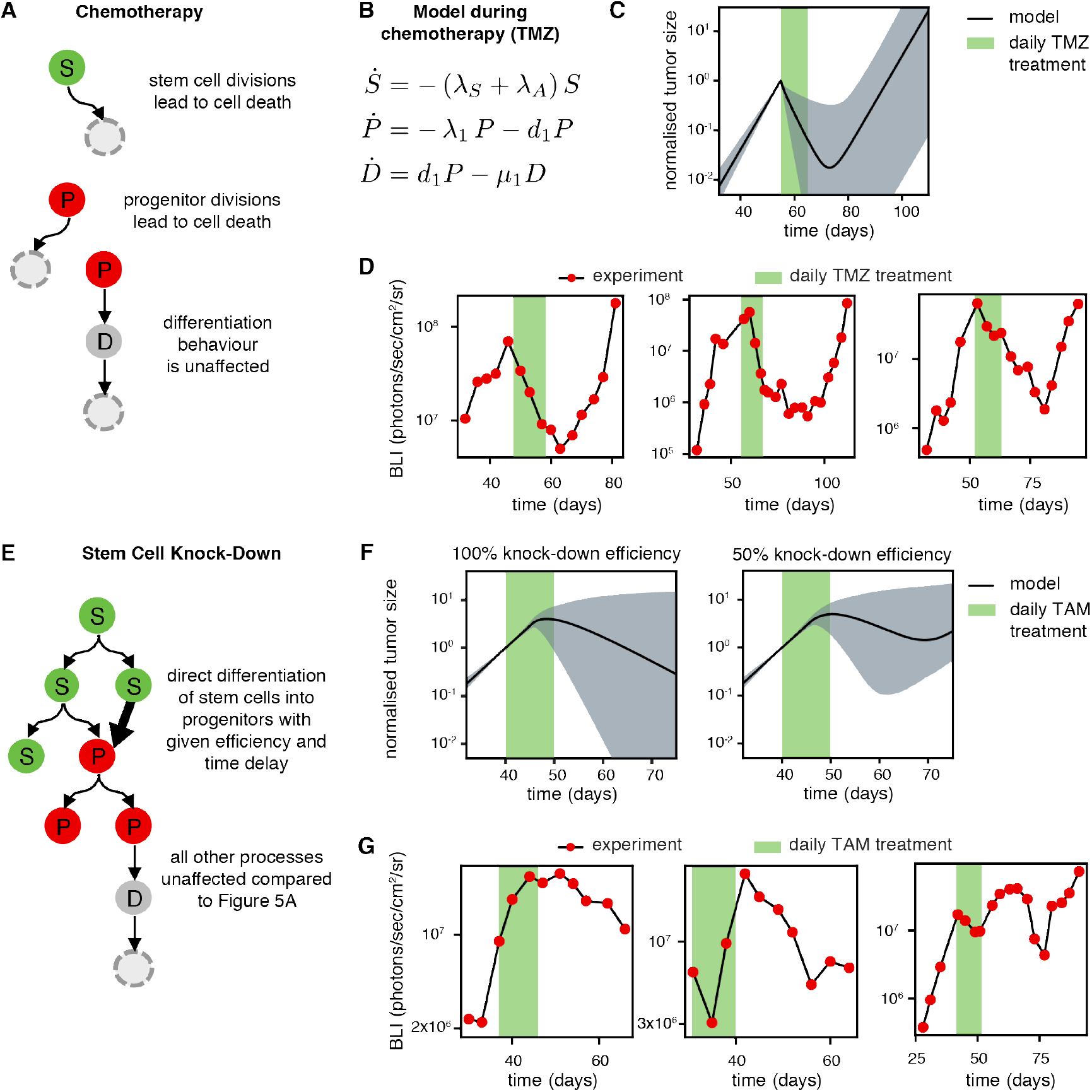
Hierarchical tumor growth model predicts treatment responses. **A**, Model from Figure 5A adjusted to incorporate effects of chemotherapeutic treatment. **B**, Translation of the model in panel A into ordinary differential equations. **C**, Model prediction of tumor size before, during and after 10 days of chemotherapeutic treatment. Shaded areas show a=0.05 posterior probability bounds. **D**, Bioluminescence imaging (BLI) of three tumor-bearing mice treated with temozolomide (TMZ) on 10 consecutive days. **E**, Model from Figure 5A adjusted to incorporate the effect of Tlx KD in BTSCs. **F**, Model prediction of tumor size before, during and after 10 days of Tlx KD in 100% (left) and 50% (right) of cells per daily drug administration. Shaded areas show α=0.05 posterior probability bounds. **G**, Bioluminescence imaging (BLI) of three tumor-bearing mice treated with tamoxifen (TAM) on 10 consecutive days to induce KD of Tlx.

As surviving BTSCs limit the effectiveness of TMZ treatment, we asked whether the alternative approach of targeting BTSCs specifically could eliminate the tumor. Tlx inactivation leads to differentiation of BTSCs [18], and we computationally simulated Tlx knockdown (KD) by turning BTSCs into progenitors with a time delay (differentiation time) of five days (Figure 6E). During the knockdown period, the tumor kept growing, but efficient Tlx KD in all BTSCs caused the tumor to regress slowly (Figure 6F, Supplemental Theory Figure T7). When we reduced the efficiency of Tlx KD to 50% (meaning that 50% of BTSCs turn into TPCs per day of the KD period), the tumor regressed for about 20 days following the end of simulated treatment but then regrew (Figure 6F). Thus, depending on its efficiency, targeting BTSCs may cause transient regression or even elimination of the tumor. To examine the response of tumors towards BTSC targeting experimentally, we modified the RGFlex system used for tracing by inserting an shRNA construct targeting Tlx into the GFP-expressing cassette (Figure S3B-D). Thus, Tlx expression was inhibited upon TAM treatment. Nestin-CreERT2;Ntva mice were used for this experiment. We found that most of the tumors kept growing during Tlx KD (Figure 6G), which is in line with both the model and our earlier data [18]. However, around 20 days after TAM injection, tumors regressed (Figure 6G). The long-term outcome of this experiment varied between individual mice. In some cases, we observed long-term stabilization of BLI signal at very low levels (Figure 6G, middle panel) while in other cases the tumor regrew (Figure 6G, right panel). These data are consistent with the model prediction and indicate varying efficiency of Tlx knockdown in the experiments. Taken together, our data show that efficient targeting of BTSCs has the potential to abrogate tumor growth.

### Molecular Signature Supports the Stem Cell Hierarchy in Mouse and Human Brain Tumors

To analyse the molecular signatures of the stem cell hierarchy in mouse GBM, and compare our results to data from human GBM samples, we performed RNA-Seq combining bulk and single-cell approaches. To differentiate between cell types, we labelled the tumor cells with RCAS-dsRed when inducing tumors with PDGFB, Akt and Fluc in Tlx-GFP;Ntv-a mice. Tlx-GFP^+^/dsRed^+^ cells (from N=4 biological replicates), containing BTSCs and early TPCs, and Tlx-GFP^−^ /dsRed^+^ cells (from N=3 biological replicates), consisting of differentiating TPCs and differentiated tumor cells, were isolated via FACS and subjected to RNA sequencing (Figure 7A).

**Figure 7.**
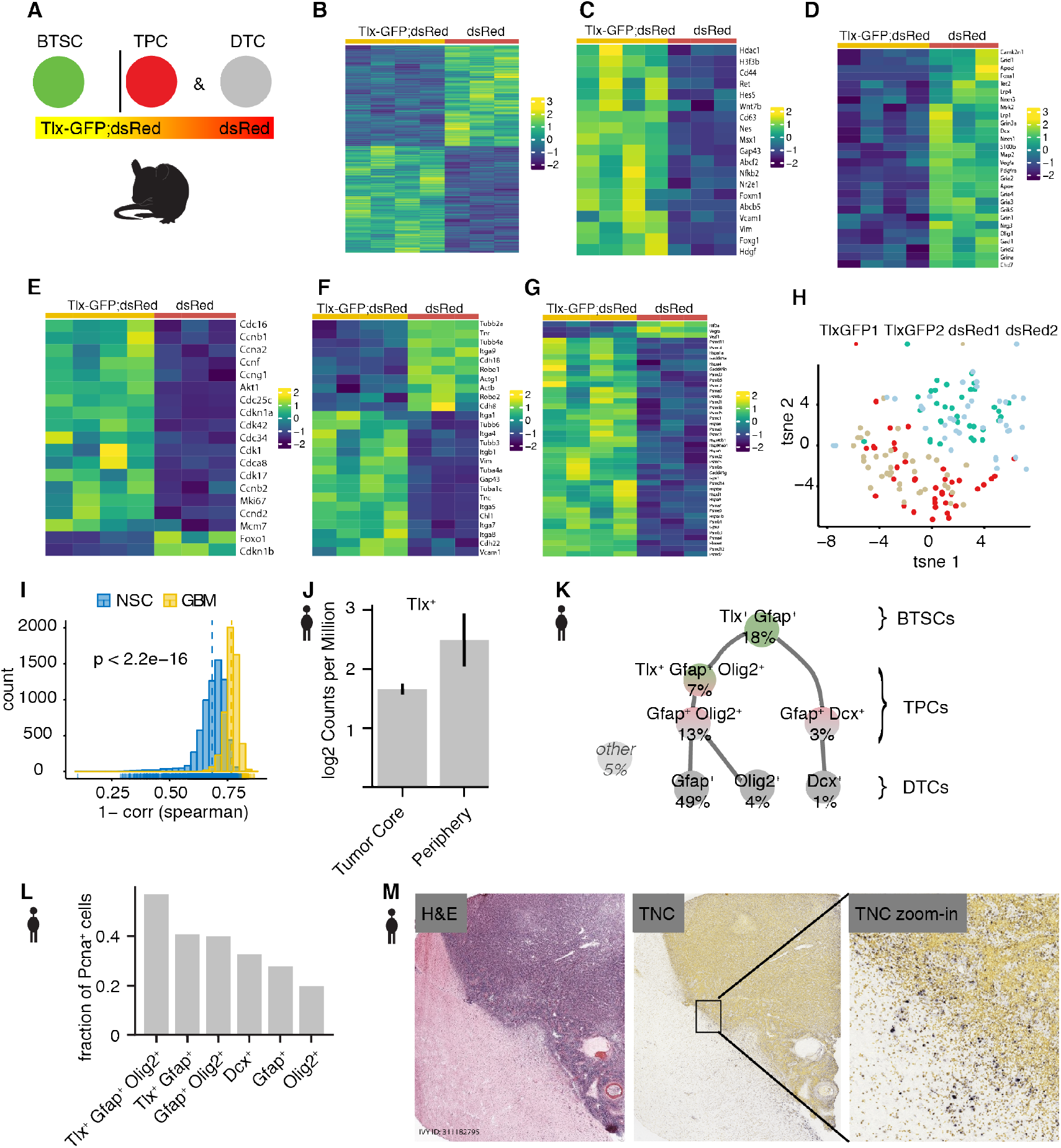
Molecular signatures of the cellular hierarchy in mouse and human glioblastoma. Panels A-I show mouse data; panels J-M show human data. **A**, Two tumor cell populations were sorted based on Tlx-GFP expression using FACS for bulk RNA-Seq. **B**, Heatmap of differentially expressed genes shows the distinct molecular signature of Tlx-GFP-positive and Tlx-GFP-negative tumor cells. All heatmaps show standardized gene expression values. N=4 mice for Tlx-GFP-positive tumor samples, N=3 mice for Tlx-GFP-negative tumor samples. **C**, Tlx-GFP positive cells are highly enriched for stem cell regulators. **D**, Tlx-GFP-negative tumor cells are enriched for neural differentiation pathways. **E**, Cell cycle genes are highly expressed in the Tlx-GFP positive tumor cells. **F**, Cell adhesion programs differ between Tlx-GFP positive and negative cells. **G**, Tlx-GFP positive cells highly express stress response genes and proteasome genes. **H**, t-SNE analysis of single-cell RNA-Seq results indicates large differences between tumors. **I**, Histogram of cell-cell distances (1 – spearman correlation coefficient) between neural stem cells and brain tumor cells respectively. The p-value is calculated using the Mann–Whitney U test. **J**, Tlx is more highly expressed in peripheral human GBM cells than in cells from the tumor core. Figure modified from www.gbmseq.org. **K**, Stem cell hierarchy of human GBM cells generated based on expression of lineage marker combinations. 5% of cells could not be unambiguously assigned to one of the groups. **L**, Percentage of cells expressing PCNA among the different populations of human GBM cells along the stem cell hierarchy in panel K. **M**, TNC expression is enriched at the tumor margin in human GBM. ISH data obtained from the IVY GBM database. H&E-Hematoxylin and eosin stain.

Indeed, Tlx-GFP^+^ and Tlx-GFP^−^ tumor cell populations had distinct molecular signatures (Figure 7B, Table S1). The Tlx-GFP^+^ population showed a stem cell signature (Figure 7C), which included known stem cell genes such as Hes5, CD44, Nestin, TNC, Vimentin and Foxg1. Gap43 expression was found to be essential for glioma cell communication via tumor microtubes [42] and was also highly enriched in the Tlx-GFP-positive cells. By contrast, differentiation programs were upregulated in Tlx-GFP^-^ cells (Figure 7D), including oligodendrocyte markers like Apod, Olig1 and Pdgfra, and astrocyte markers such as Apoe and S100b. We also observed neuronal transcripts, such as Map2, Dcx, Chd7 and mRNAs encoding gamma-aminobutyric acid and glutamate receptors. Thus, the Tlx-GFP^-^ tumor fraction is enriched in differentiated tumor cells.

Further, we found that the GFP^+^ cells were enriched for cell-cycle genes, including several cyclins (Figure 7E). This finding provides molecular underpinning to our observation that active BTSCs drive tumor growth while progressive differentiation of tumor cells is accompanied by loss of proliferation (Figure 5A). Tlx-GFP^+^ and Tlx-GFP^-^ cells also expressed distinct sets of cell adhesion molecules (Figure 7F). In particular, Tlx-GFP^+^ cells expressed several integrins (Itga4, Itga5, Itga7, Itga8 and Itgb1), which may support the migration of BTSCs. Of note, the Tlx-GFP^+^ population was highly enriched for heat shock stress response and protein quality control pathways. This set of genes also includes 22 genes encoding different proteasome subunits (Figure 7G), which are known to be essential for the degradation of stress-induced misfolded proteins [43].

Taken together, these data confirm that GBM tissue retains signatures of the physiological stem cell hierarchy, with active BTSCs giving rise to more differentiated tumor cells. At the same time, we uncover distinct tumor-cell-specific features, including the high proliferation of BTSCs and early TPCs (consistent with Figure 5A), the support of BTSC migration by integrin expression (in agreement with Figure 1), and the upregulation of stress response pathways that may protect BTSCs during cytotoxic treatment. Using human GBM data from cBioPortal [44,45], we asked whether genes enriched in mouse BTSCs have prognostic value in the human context. Indeed, we found that high expression of these genes indicates poor survival prospects (Figure S4). This in turn may imply that a high frequency of BTSCs in human GBM correlates with poor survival as has been suggested for cancer stem cells in general [46].

Next, we analyzed RNA-Seq results from single cells stemming from two separate tumors. Tlx-GFP^+^/dsRed^+^ and Tlx-GFP^-^/dsRed^+^ cells were sorted via FACS in the same way as for the bulk RNA-seq (Figure 7A). When subjected to t-distributed stochastic neighbor embedding (tSNE), cells from different tumors clustered together, irrespective of whether they were Tlx-GFP-positive or -negative (Figure 7H). This suggests that intertumoral heterogeneity dominates over intratumoral heterogeneity, which has also been reported for single-cell RNA-Seq data of human tumors [47]. Next, we examined the correlation (Spearman correlation coefficient) between the tumor cells, using RNA-Seq results from normal Tlx-positive neural stem cells as control. The overall distance in gene expression between brain tumor cells was significantly larger than that between normal neural stem cells, thus corroborating the large molecular heterogeneity of mouse tumor cells (Figure 7I).

Further, we compared the finding of our murine transcriptome analyses to published single-cell RNA-Seq data of human brain tumors. Consistent with our findings on tumor cell heterogeneity, the human data showed no clear subgroups under unsupervised tSNE analysis of (Figure S5A) [47–50]. Moreover, in three of the studies we considered, Tlx was either not detected at all or at a low level in few cells [47,49,50], likely because of insufficient sequencing depth. In the fourth study, which separately analyzed GBM core and periphery, Tlx was captured more frequently (Figure S5B) [48]. Notably, Tlx expression was enriched in the periphery (Figure 7J), which is in line with our finding that Tlx-positive cells are located preferentially in the invasive front of mouse GBMs.

We applied our mouse-derived knowledge of the cellular hierarchy and divided the human tumor cells into the following subtypes: Tlx/Gfap doublepositive cells likely representing BTSCs, Tlx/Gfap/Olig2 or Tlx/Gfap/Dcx positive cells being putative early TPCs, and Tlx-negative but Gfap/Olig2 positive cells, which are TPCs with bipotent glial differentiation potential. Tlx-negative cells expressing either GFAP, Olig2 or Dcx were considered differentiated tumor cells. When arranging these subtypes into a cellular hierarchy (Figure 7K) we found that the resulting tumor composition with 18% BTSCs, 23% TPCs and 54% DTCs (the remaining 5% could not be assigned clearly) agreed with our murine GBM data (Figure 5F). Finally, we asked whether the human tumor cells can also be separated by their proliferative activity. Considering the cell proliferation marker PCNA we found that early TPCs (Tlx/Gfap/Olig2) had the highest proliferation index, followed by BTSCs and late TPC (Tlx^-^/Gfap/Olig2) and, finally, DTCs (Figure 7L). Using the IVY database [51], we found that the expression of Tenascin C (TNC), which is enriched in mouse BTSCs (Figure 7F), occurs preferentially at the boarder of human GBM (Figure 7M), further supporting that our mouse-based findings are also relevant for human tumors.

## Discussion

In this study, we combined single-cell analysis and bulk tumor imaging on the same mouse model, integrated the data into a mathematical model of tumor growth and quantified proliferation and differentiation rates of tumor stem and progenitor cells. We obtained four main findings: First, the majority of GBM grew exponentially until mice had to be sacrificed, indicating a lack of growth limitations by nutrients or space constraints. Second, the tumors are hierarchically organized, containing BTSCs with self-renewal capacity and non-BTSCs that are short-lived in vivo and show no detectable dedifferentiation during tumor expansion. Third, the bulk tumor growth rate is determined by actively proliferating BTSCs. Fourth, to maintain exponential growth, BTSCs must migrate outwards. In addition, we provide a transcriptional analysis supporting the mechanisms identified using the mathematical model and demonstrate the transferability of our findings to human GBM.

Exponential brain tumor growth has also been reported for human patients: A study on human low-grade glioma patients with at least eight serial MRI time points per patient finds exponentially increasing tumor volumes [52]. In another study, growth data of untreated glioblastomas could be fitted best by a Gompertzian description (although this relies on only two time points per patient and no penalty was applied for the larger parameter count of the Gompertz growth law over the simpler exponential model) [30]. Here, we observed signs of growth limitations of Gompertzian type during the lifetime of about one fifth (22%) of the studied mice. This variability in growth laws may be caused by the exact brain region the tumor is occupying as well as by differences in absolute size.

Further, exponential growth implies the absence of space constraints for BTSC division, which we show here to be overcome by outward migration of BTSCs. Invasive and infiltrative growth is a major hallmark of human GBM, and mathematical models have previously highlighted the importance of tumor cell migration for fast tumor expansion [3,31,53,54]. The migration of BTSCs also suggests that the niche for BTSCs is constantly changing as the tumor grows. This may explain why stress response and protein quality pathways are upregulated in BTSCs. Several previous reports suggested a hypoxia niche for BTSCs supposedly at the core of GBM [55]. Our study reveals a novel environment for BTSCs, which calls for additional studies dissecting the heterogeneity of BTSCs and their niches in vivo. The tumor margin-associated relapse [35] and the enrichment of BTSC markers at the boarder which we observed in human GBM patients is consistent with our murine observations.

At the molecular level, we found that BTSCs highly express heat shock and proteasome proteins which are involved in stress-related protein quality control, suggesting that BTSCs may be vulnerable to proteasome inhibitor treatment. Indeed, this direction is currently being investigated in a phase III clinical trial against human GBM with the second-generation proteasome inhibitor Marizomib [43] (https://clinicaltrials.gov/ct2/show/NCT03345095). Moreover, our findings point to BTSC migration as a potential therapeutic target meriting further study.

Using observations and measurements from several mouse experiments, we have assembled and quantified a hierarchical model of brain tumor growth (Fig. 5A). Our model does not involve dedifferentiation of Tlx-GFP-negative cells into Tlx-GFP-positive BTSCs during tumor progression as we were unable to observe this experimentally. This result opposes the notion of cellular dedifferentiation in tumors [12] and leaves symmetric BTSC divisions as the driving force of tumor growth. Indeed, we found that the symmetric division rate of BTSCs determined from cell proliferation assays in vivo fully accounts for the growth rate of the bulk tumor obtained by bioluminescence measurements (average doubling time of approximately 3.5 days), providing a quantitative match between cellular and macroscopic tumor properties.

Our rate estimates concerning BTSC behavior describe the average dynamics of this population. Importantly, they do not exclude the existence of a quiescent subpopulation or phase. Here, based on our quantified model, we find that stem cells make up around 19% of the tumor mass while proliferating progenitors and differentiated cells contribute 24% and 57% respectively (Figure 5C). Since the proportion of stem cells is directly linked to tumor aggressiveness [46], this high stem cell fraction may explain poor glioblastoma survival. This is supported by our finding that genes which are found to be highly expressed in mouse BTSCs can serve as predictors for poor survival in human GBM patients.

In summary, we provide a systematic multi-scale analysis of glioblastoma growth in vivo which involves molecular, single-cell and bulk experiments and integrates their results using spatio-temporal models. Our approach allowed us to dissect the individual contributions of GBM subpopulations to tumor progression and to identify prospective targets for GBM treatment.

## Supporting information

Supplementary Theory

## Acknowledgments

We would like to thank the DKFZ preclinical cancer center for assistance for animal experiments; the Imaging and Cytometry, Genomics and Proteomics Core Facilities of the DKFZ and the Carl Zeiss Imaging Center in the DKFZ for their support. This work was supported by the Helmholtz Association (VH-NG-702), the Deutsche Krebshilfe (110226 to H-KL), the ERC (European Research Council) consolidator grant (647055) (to H-KL) and the BMBF Sys-Glio consortium (H-KL and TH).

## Author contributions

H-KL conceived and supervised the project. MAK performed clonal tracing and RNA-Seq experiments. LB generated the mathematical models with help from CB. TH supervised the mathematical modeling. YZ performed the clonal and retroviral tracing experiments. PZ performed the mirFlex experiment. CS performed bioinformatics analysis. WF, MQ, GB, AAA, AN, ZZ, KH and MT helped with experiments. HA, KH and OF provided technical assistance. H-KL, TH and LB wrote the manuscript. All authors contributed to the editing of the manuscript.

## Material and Methods

### Animal Experiments

Mice were housed according to international standard conditions and all animal experiments complied with local and international guidelines for the use of experimental animals. For survival analysis, animals were sacrificed once reaching the endpoint based on criteria approved in the animal procedure. The Ntv-a mouse was kindly provided by Eric Holland. The Tlx-CreER^T2^ animal was generated as described [26]. The Tlx-GFP reporter animal was described before [56]. Confetti mice were from Jackson Laboratory. Following transgenic mice lines were used for experiments Tlx-GFP/ N-tva, Tlx-CreER^T2^/ Ntv-a /R26-Confetti, Nestin-CreER^T2^/Tlx^flox/flox^, Tlx-CreER^T2^/Tlx-GFP/ Ntv-a /R26-Confetti, Ntv-a;NestinCreER^T2^, which were described before [18].

### Primary Brain Tumor Induction

DF-1 cells were used to produce and maintain RCAS viruses. The cells were maintained in DMEM (ATCC, 30-2002) supplemented with 10% FBS (ATCC, 30-2020) and 1% Penicillin-Streptomycin (Life Technologies, 15140122) in a humidified incubator with 5% CO2 at 39 °C, and split every two days. For transfection, early passage of cryopreserved DF-1 cells was thawed, and after expanding, the cells were plated into 6-well plates so that on the day of transfection, the confluence of cells should reach 50-70%. 1.6 μg DNA and 4 μl FuGENE HD transfection reagent (Promega, E231A) was used for 6-well format transfection according to the manual. During the passage of these cells, it is very important to maintain cells producing virus carrying different transgenes separately.

RCAS virus can specifically infect the cells expressing TVA receptor. Our strategy of inducing brain tumor is to inject DF-1 cells producing RCAS virus carrying oncogenes into the subventricular zone (SVZ) of the brain in Nestin-TVA mouse. RCAS-AKT, RCAS-PDGFB and RCAS-luciferase vectors were transfected into DF-1 cells respectively according to the procedure described above, then the cells were split regularly every two days. Since DF-1 cells are permissive for viral infection and replication, after 3-4 passages most of the cells should be infected and are able to produce a high titer of RCAS virus. On the day of injection, make single cell suspension for each of the DF-1 cell types and measure the concentration, mix different cell types in a way where each of cell types can get a final concentration of 4 × 10^4^ cells/μl in the mixture. In the case of injection together with RCAS-miRFlex/RGFlex, the final concentration of cells transfected with RCAS-miRFlex/RGFlex was 7 × 10^4^ cells/μl and that of the other three plasmids was 3 × 10^4^ cells/μl. 1μl of the cell mixture was injected into the lateral ventricle in the left brain hemisphere of neonatal Nestin-TVA animals using a 10 μl Hamilton syringe. The injection point locates at 1-2 mm left lateral of bregma point, after penetration of the skull, the needle is inserted around 1.5 mm before injecting the cell suspension.

### Retrovirus Packaging and Injection

GP2-293 cells stably expressing gag and pol viral genes were transfected with CAG-nls-dsRed-IRES-dsRed and pVSV-G via Lipofectamine reagent. A two-day incubation was left for viral production before collection. The supernatant was collected into 50 ml Falcon tubes and centrifuged for 10 min at 3000 rpm to pellet cells to avoid blocking during filtration. The supernatant was then filtered (0.45 μm) into the ultracentrifuge tube and was centrifuged at 25,000 rpm at 4 °C for 2 h. The supernatant was removed and 2 ml Opti-MEM was applied for resuspension followed by the second ultracentrifugation at 25,000 rpm at 4 °C for 2 h. The supernatant was removed as much as possible and 80 μl sterile PBS was applied for resuspension. The tube was then left on ice for shaking for 1 h. The entire resuspended liquid was transferred to an Eppendorf tube. To maximize retrovirus yield, additional 20 μl sterile PBS was added to the ultracentrifuge tube to collect leftover viruses and was then pooled with the 80 μl resuspension. Ten-microliter aliquots were stored at −80 °C.

Retrovirus injection was conducted via stereotaxic surgery (KOPF®). Animals were group housed and were kept under a 12 h light/dark cycle. Mice were anesthetized with a mixture of ketamine (100 mg/kg body weight) and xylazine (10 mg/kg body weight). According to bio-luminescence images, the coordinates of the tumor were generated for injection guidance. Retrovirus (1.5 μl) was injected into the region with the highest bio-luminescence signal.

### TMZ Administration

Temozolomide (TMZ) was dissolved in DMSO to prepare 25 mg/ml stock solution. Before injection, the stock solution was further diluted in saline (0.9% NaCl) to make 5mg/ml working solution. When the tumor volume reached the value of 1-5 × 10^7^ photons/sec based on the BLI, TMZ was intraperitoneally administrated to animals at a dosage of 20 μl/g body weight, once per day for 10 days.

### Tamoxifen Administration

In the experiments for lineage tracing in Tlx-CreERT^2^/Nestin-TVA/R26-Confetti and Tlx-CreER^T2^/Tlx-GFP/Nestin-TVA/R26-Confetti, when the tumor volume reached the value of 1 × 10^6^ photons/sec based on the BLI, 1 injection of tamoxifen (250 μg/g body weight) was administrated to induce recombination in Tlx expressing cells, so that both Tlx expressing cells and all their progenies (daughter cells) would be labeled by one of the four fluorescent colors, blue, green, red or yellow permanently. With this protocol, we did not observe differently colored cells in the same area, which suggests we achieved very sparse labeling of single Tlx^+^ cells. The mice were analyzed 5, 10, 20, 26 or 37 days after the administration of tamoxifen injection for analysis.

In the experiments of Tlx knockdown and RGFlex tracing, when the tumor volume reached the value of 1 × 10^6^ photons/sec based on BLI, 100 μl

Tamoxifen (10 mg/ml in sunflower seed oil with 10% ethanol) was intraperitoneally administrated to animals, once per day for 10 days. For the Tlx-KO and TMZ experiment, animals received TMZ first, and TAM was given after finishing TMZ experiment. The survival results of Tlx-KO group were described previously [18].

### Bioluminescence Imaging

Since DF-1 cells producing RCAS-firefly luciferase were injected as well, the luciferase transgene will be taken up together with oncogenes by Nestin-expressing cells and carried by tumor cells during the tumor progression, which facilitates us to monitor the tumor growth by bioluminescence imaging. The imaging started when the animals were 4 weeks old, and continued afterward with a frequency of twice every week. Before the imaging, animals were intraperitoneally given D-luciferin (Biocat, 7903-1G-BV, dissolved in PBS to make 30 mg/ml working solution), the substrate of firefly luciferase, at the dosage of 5 μl/g according to the body weight of the animals. The animals were anesthetized by isoflurane using the XGI-8 system. 10 min after D-luciferin administration, the animals were imaged using IVIS bioluminescence imaging (BLI) system. The parameters for imaging were set as 60 s exposure time; medium binning and 6 images were taken each after 60 s.

### Immunofluorescence Staining

Mice were euthanized, followed by whole body perfusion and fixation with 4% PFA. The fixed brain with the tumor was isolated and further fixed overnight at 4 °C. Next day, the brains were transferred to 30% sucrose in PBS and kept at 4 °C. Fixed brains were cut to 40-60 μm floating sections and kept at−20 °C in cryoprotectant solution. The sections were washed in PBS, subjected to Hydrochloric Acid (HCl)-based antigen retrieval if thymidine analog staining was intended, blocked with 5% normal swine serum in PBST (0.2% Triton X100 in PBS), and then incubated with primary antibody overnight at 4 °C. On the second day, the sections were washed with PBS to get rid of residual primary antibody and incubated with secondary antibodies for two hours in the dark at room temperature. Secondary antibodies conjugated with different Alexa fluorophore (Life Technologies) were chosen depending on their reactivity to the species of primary antibody. Dilution of 1:400 was used for all the Alexa fluorophore-conjugated secondary antibodies. After incubation with secondary antibody, the sections were washed with PBST with or without DAPI and finally subjected to mounting with VECTASHIELD Mounting Medium.

### Imaging

Fluorescent images were taken using Zeiss LSM780 confocal microscopy. The emission spectrum of each fluorescent protein (including RFP, GFP and YFP) and fluorophore including DAPI, Alexa Fluor405, Alexa Fluor488, Alexa Fluor555, Alexa Fluor594, Alexa Fluor633, was captured in samples with corresponding individual staining, and saved in spectrum database of the system. Imaging protocol with a different combination of these spectrums under online fingerprinting mode was used depending on the fluorescent protein or secondary antibodies used for each staining. Processing of the images and quantification of cell numbers for each cell type was conducted manually.

For clonal tracing of confetti mice, 60 μm sections from confetti mice were mounted on slides using VECTASHIELD Mounting Medium without DAPI and kept at 4 °C before microscopic analysis. Mouse sections were numbered consecutively and imaged in the same order, clone size was determined by counting all the cells found to be on the same area on adjacent sections.

### Brainbow Clonal Analysis

For counting of Brainbow clones, ten to twelve consecutive free-floating sections from each animal were selected and mounted onto a glass slide with VECTASHIELD Mounting Medium without DAPI. The sections were snake-scanned through the eyepiece, with constant changing of filters to search for cells of different Brainbow colors. Once those cells were identified, they were documented by confocal scanning described in the imaging section. Cells from different sections were considered belonging to the same clone if they share the same Brainbow color and appear at a similar location.

For 3D reconstruction of Brainbow cell clones, 12 consecutive free-floating sections (60 μm) from each animal were scanned with a 20X objective on Zeiss AxioScan as z-stacks with a 10-μm interval. The resultant micrographs were maximal-intensity-projected and exported from ZEN and then manually aligned in GIMP (GNU Image Manipulation Program) to recapitulate clonal location in 3D. These images were assembled as an image stack in ImageJ/FIJI. RFP^+^ cells were identified in the red channel, while nls-GFP^+^ and YFP^+^ cells were distinguished by their sub-cellular localization in the green channel. Cell count and location were documented with the “Cell Counter” plugin in ImageJ/FIJI. The plots for the XY-plane were generated in R with the ggplot2 package.

### MicroRNA Design and Efficiency Test

Four miRs designed for targeting TLX and a negative control miR (BLOCK-iT^™^ RNAi Designer, ThermoFisher) were cloned into pcDNA3.1 and cotransfected with a vector expressing mouse TLX into HEK293T cells. Forty-eight hours later, the cells were harvested for TLX expression analysis with Western blot. The miR with the highest knockdown efficiency was then cloned into RCAS vector for in vivo experiments.

### Western Blotting and Analysis

The whole protein extraction procedure was performed on ice or in 4 °C environment. Cells were washed with PBS to remove the residual medium. 100 μl of total protein extraction buffer was added into the wells of 6-well plate. Cells were scraped off and transferred into 1.5 ml Eppendorf tubes. The cell suspension was kept on ice for lysis for 5 min. 10 μl 10% NP40 was added to the suspension and the tube was vortexed for 10 s. Then add 8μl 5M NaCl and rotate the lysate for 15 min at 4 °C. After that, the lysate was centrifuged at 5000 g for 5 min, and the supernatant was collected. The protein concentration was measured using Bradford method. After mixing with 6X protein loading dye, the protein solution was boiled at 95 °C for 5min. Then sonicate the protein solution using 10 cycles of 15 s on-off program. After that, the protein solution was ready for western blot.

Equal amount of protein was loaded into SDS-PAGE gels. The samples were first run in 5% stacking gel at 80V for 30min, then separated in 10% gel at 120 V for 90 min using Bio-Rad Mini Trans-Blot cell. Then the protein was transferred onto Nitrocellulose membrane at 15 V for 45 min using Bio-Rad Trans-Blot Semi-Dry transfer cell. The blot was blocked in 5% skim milk for 1 h at room temperature, and then incubated with primary antibody overnight at 4 °C, followed by incubation with HRP-conjugated secondary antibody for 2 h at room temperature. Dilution of 1:1000 for rabbit anti-TLX, 1:1000 for rabbit anti-β-tubulin (Cell signaling, 2128s), 1:1000 for chicken anti-GFP (aves, GFP-1020), and 1:10000 for all the secondary antibodies were used. After incubating together with the chemiluminescent HRP substrate (Millipore, WBKLS0500), the blots were developed using LAS-3000 plus system. Intensity of the bands were measured in Image Lab^™^ (Bio-Rad).

### FACS (both single cells and Bulk cells FACS)

Tumors were induced in Tlx-GFP/Ntv-a mice using RCAS-AKT, RCAS-PDGFB and RCAS-Fluc RFP retroviruses. Brain tumor tissues were isolated in Phosphate buffered saline solution (PBS) and enzymatically dissociated with 0.05% Trypsin/EDTA (Gibco) in HBSS containing 2 mM glucose at 37 °C for 30 min. During incubation, the tissue was repeatedly triturated with a fire-polished Pasteur pipette. Enzyme activity was stopped by addition of equal volume of 4% BSA in Earle’s Balanced Salt Solution (EBSS, Gibco). The cell suspension was filtered through a 70 μm cell strainer and centrifuged at 1200 rpm for 5 min and the pellet was re-suspended in 0.9 M sucrose in 0.5 x HBSS (Gibco). After further centrifugation for 20 min at 2000 rpm, the cell pellet was re-suspended in 2 ml of 4 % BSA in EBSS solution and placed on top of 12 ml of 4% BSA in EBSS solution, centrifuged again for 9 min at 1500 rpm. The resulting pellet was then suspended in PBS containing Recombinant RNase Inhibitor, 1 unit/μl reaction (Clontech). After incubation with PI (1:1000) for 2 min; either the single cells or 1000 cells per tube were sorted at a FACS Aria (BD). Two population of cells were sorted, RFP positive and RFP / Tlx-GFP positive cells.

### RNA Isolation from Bulk Cells

RNA extraction using the Arcturus PicoPure RNA Isolation kit was performed according to the manufacturer’s protocol (Life technologies). Briefly, 100μl of extraction buffer was added to the cells, followed by brief incubation at room temperature. While the lysate was incubated, the purification column was wetted (Pre-condition) by adding 250*μ*l of condition buffer, followed by centrifugation at 16,000 × g for 1 min. 100 *μ*l of 70% ethanol was then added to the lysate, mixed well and the whole mixture was transferred to the prepared purification columns. Purification columns were centrifuged for 2 min at 100 × g, followed by centrifuging at 16,000 × g for 30 s to remove flow through. 100 *μ*l of wash buffer 1 was added to the purification columns, which were then centrifuged for 1 min at 8,000 × g. DNase treatment may be performed directly within the purification column. 10 *μ*l DNase I stock solution (Qiagen, catalog#79254) was mixed with 30 *μ*L Buffer RDD. 40 *μ*L DNase incubation mix was applied directly into the purification column membrane followed by Incubation at room temperature for 15 min. 40 *μ*l of wash buffer 1 was added to the purification columns, which were then centrifuged for 15 seconds at 8,000 × g. After that, 100 *μ*l of wash buffer 2 was added to the purification columns, which were then centrifuged for 1 min at 8,000 × g. Another 100 *μ*l of wash buffer 2 was added to the purification columns. Purification columns were centrifuged at 16,000 × g for 2 min. Another spin at 16,000 × g for 1 min was performed to remove the wash residue in the columns. Purification columns were transferred to new Eppendorf tubes and 11 *μ*l of elution buffer was carefully added onto the column membrane followed by 1 min incubation at room temperature. Purification Columns were centrifuged at 1,000 × g for 1 min, and subsequently at 16,000 × g for 1 min to elute RNA. 10 *μ*l of RNA solution was obtained.

### RNA Sequencing and Bioinformatic Analysis

For single cell RNA-Seq, RT-PCR and cDNA synthesis were performed using SMARTer Ultra Low Input RNA Kit v3 following manufacturer’s protocol (Clontech). Single cells were used directly as starting material without RNA extraction to avoid extra loss of RNA. First strand cDNA was synthesized using 3’ SMART CDS Primer II A and SMARTer IIA Oligonucleotide with SMARTScribe Reverse Transcriptase. dT priming was used to eliminate DNA contamination. The cDNA was then amplified by 23 cycles of LD PCR and purified using Agencourt AMPure XP kit. Amplified cDNA was validated using Agilent 2100 BioAnalyzer.

For bulk cell RNA-Seq, RT-PCR and cDNA synthesis were performed using SMARTer Ultra Low Input RNA Kit v3 following manufacturer’s protocol (Clontech). RNA isolated from 1000 cells was used as starting material. First strand cDNA was synthesized using 3’ SMART CDS Primer II A and SMARTer IIA Oligonucleotide with SMARTScribe Reverse Transcriptase. dT priming was used to eliminate DNA contamination. The cDNA was then amplified by 18 cycles of LD PCR and purified using Agencourt AMPure XP kit. Amplified cDNA was validated using Agilent 2100 BioAnalyzer.

Libraries were constructed using the Nextera XT kit as recommended by the manufacturer (Illumina, San Diego, USA). 0.5 ng of cDNA was tagmented, utilizing 5 μl of Amplicon Tagment Mix with 10 μl Tagment DNA Buffer. Tagmentation reaction was performed by incubation for 20 min at 55 °C. Next, index primers were added together with Nextera PCR Master Mix. Tagmented cDNA was amplified by limited-cycle PCR (10 cycles). Libraries were purified using AMPure beads (Beckman Coulter, High Wycombe, UK) and quality validated using Agilent 2100 BioAnalyzer. Libraries were normalized and 8 nM pooled libraries were then sequenced on Illumina HiSeq2000.

For all single-cell/bulk libraries from neural stem cells or brain tumors, we obtained the gene expression profiles via mapping the sequencing reads to mouse mm10 transcriptome by RSEM (v1.3.0) [57]. For single cells from brain tumors, we kept cells with 200 to 3500 genes and less than 10% of reads on mitochondrial genes and projected the cells into low dimension space using Monocle [58,59] in R. We calculated the Spearman correlation coefficient for single cells in neural stem cell populations and brain tumor populations using log2 transformed TPM values.

We used the R package DESeq2 [60] to estimate the differentially expressed genes between bulk Tlx-GFP^+^/dsRed^+^ and Tlx-GFP^-^/dsRed^+^ using the estimated counts from RSEM results. The statistically significant genes were selected with fdr <= 0.1. Heatmaps for selected genes were produced with ComplexHeatmap [61].

We downloaded the raw single-cell RNA-seq data of GSE84465 from GEO and estimated the gene expression profiles via kallisto [62]. Cells having 1000 to 8000 genes were used in the downstream analysis. Neoplastic cells are selected according to the accompanying publication. We used the threshold log2(TPM+1) > 1 to identify cells expressing marker genes.

### Mathematical Modelling and Code Availability

All code related to mathematical models, simulations and parameter estimation can be found at https://github.com/LiBuchauer/exponential_GBM. The computations were performed using python 3.5 and relying on the following external packages: numpy [63], matplotlib [64], seaborn [65], lmfit [66], pandas [67], scipy [68], emcee [69], corner [70] and Cython [71]. Detailed explanations regarding the models are available in the *Supplementary Theory*.

## Supplementary Figures

**Figure S1.**
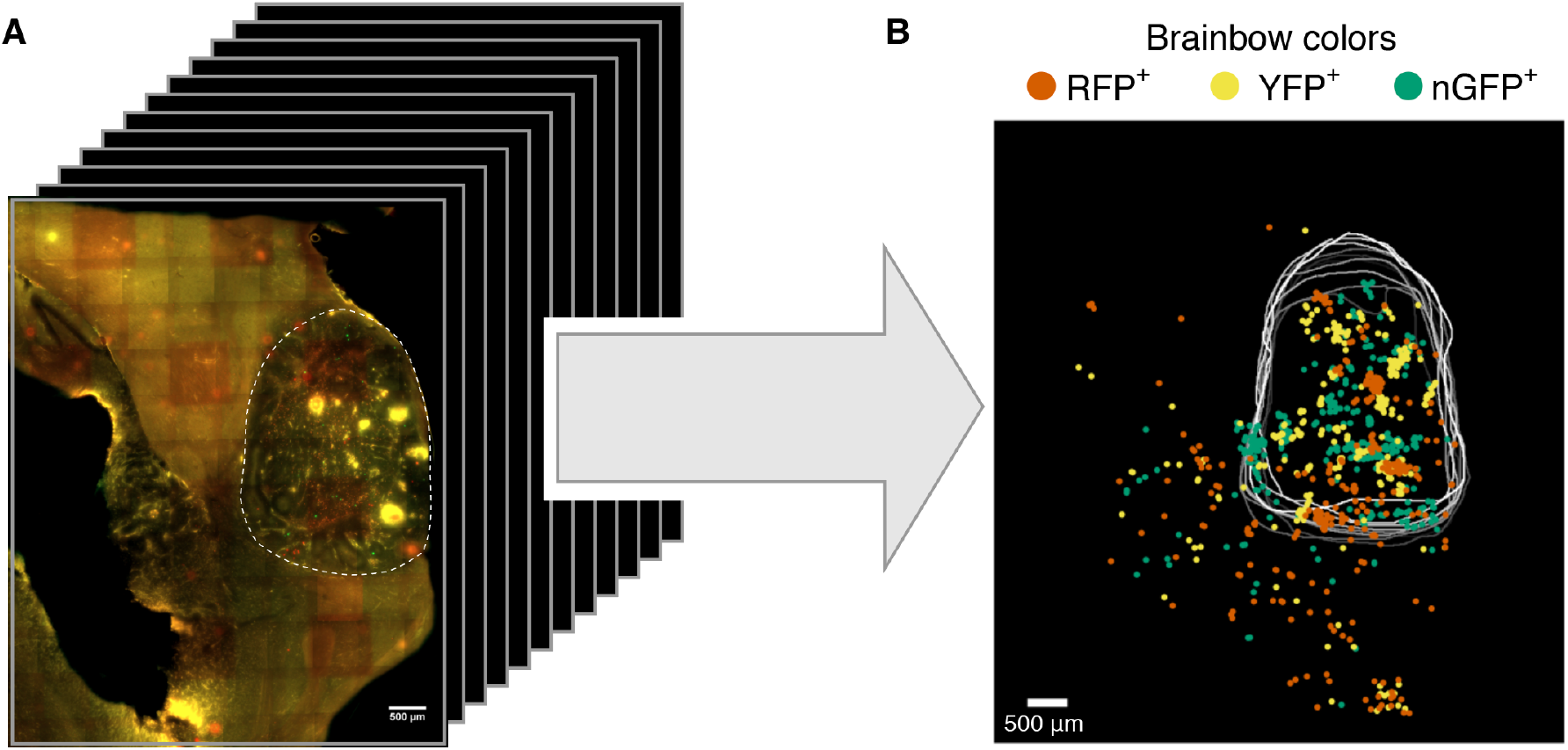
Three-dimensional reconstruction of multiple BTSC-derived clones. **A**, Twelve consecutive sections were scanned and aligned manually. The tumor was outlined with a white dashed line. Scale bar, 500 μm. **B**, Cells with different Brainbow colors were identified and plotted across the sections showing their relative positions in the tumor, outlined with white lines.

**Figure S2.**
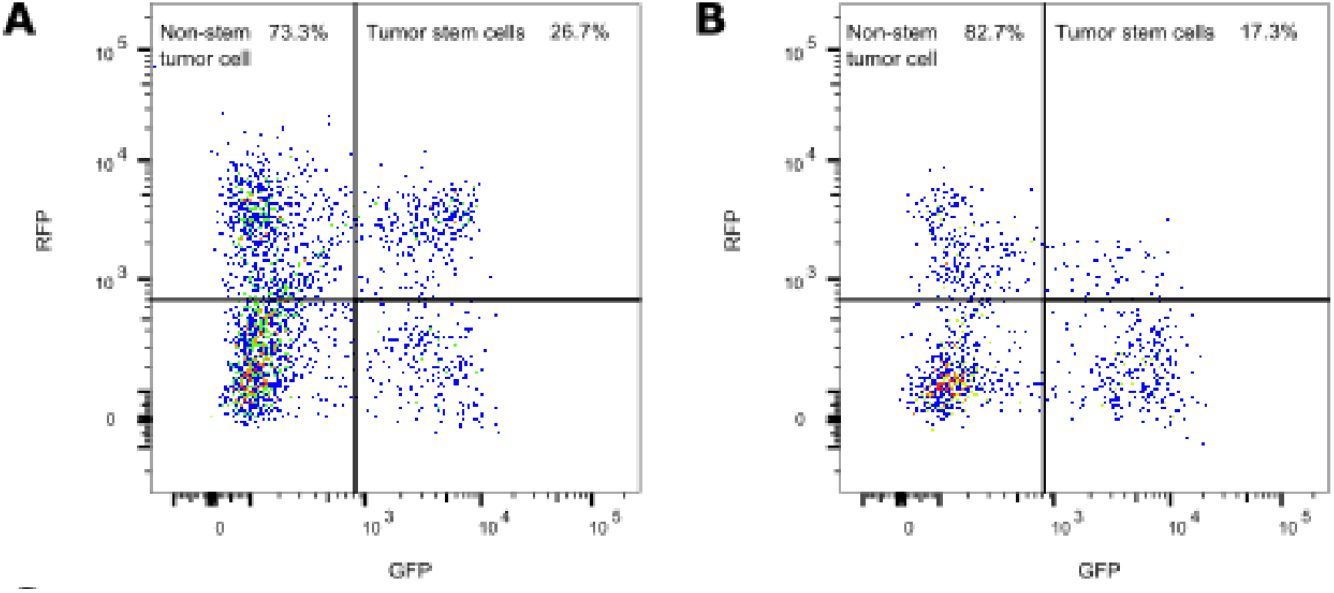
Percentage of Tlx expressing Cells in RFP-labelled tumors. **A** & **B**, FACs analysis yielding the percentage of Tlx-GFP;RFP double positive cells among all RFP-positive tumor cells in two biological replicates.

**Figure S3.**
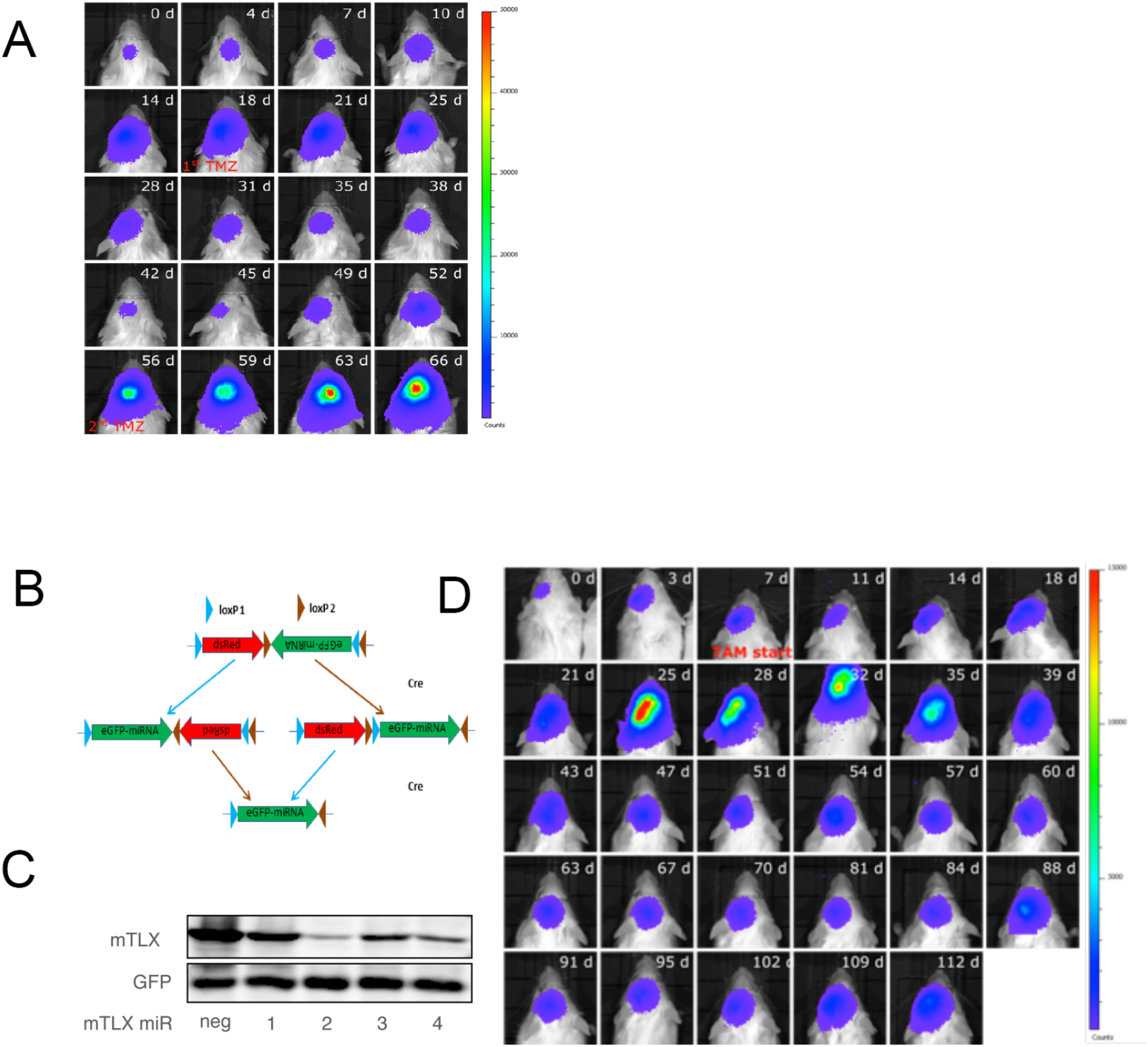
Tumor response towards treatment. **A**, BLI imaging of tumor response to TMZ treatment in an exemplary animal. **B**, Schematic illustration of an inducible miRNA-based knock-down system. **C**, Four microRNAs and a negative control were delivered to mouse Tlx-transfected HEK293 cells to test Tlx knockdown efficiency. A Western blotting demonstrated the extent of knockdown by these microRNAs with GFP as an internal control. **D**, BLI imaging of tumor response to Tlx inducible knock down upon TAM treatment.

**Figure S4.**
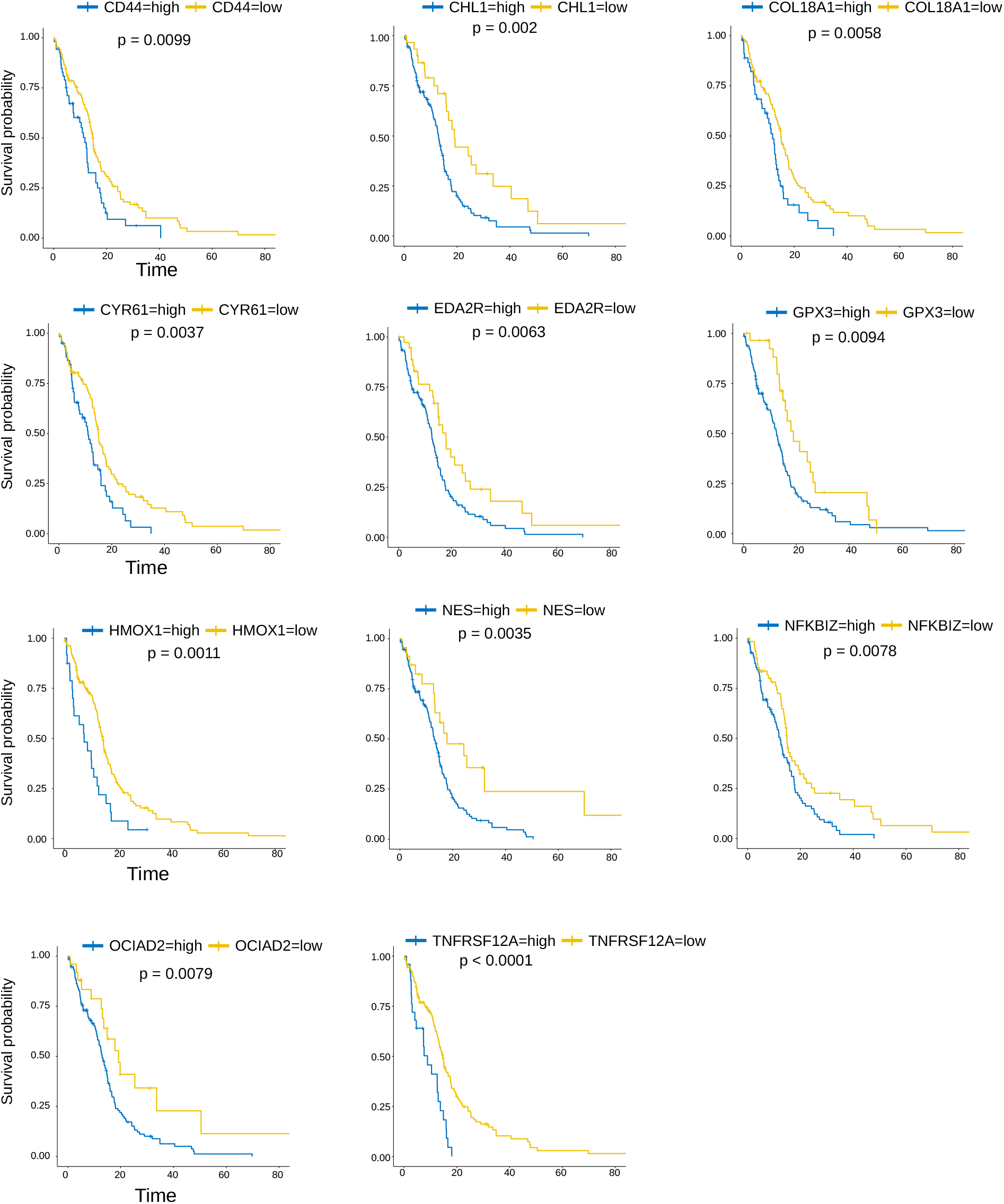
Stem Cell Signature and Patient Prognosis. Highly enriched genes of mouse BTSCs were used to analyze their prognostic values in human GBM patients. Results from the TCGA (cBioPortal) database suggest high expression of BTSC enriched genes correlates with poor survival.

**Figure S5.**
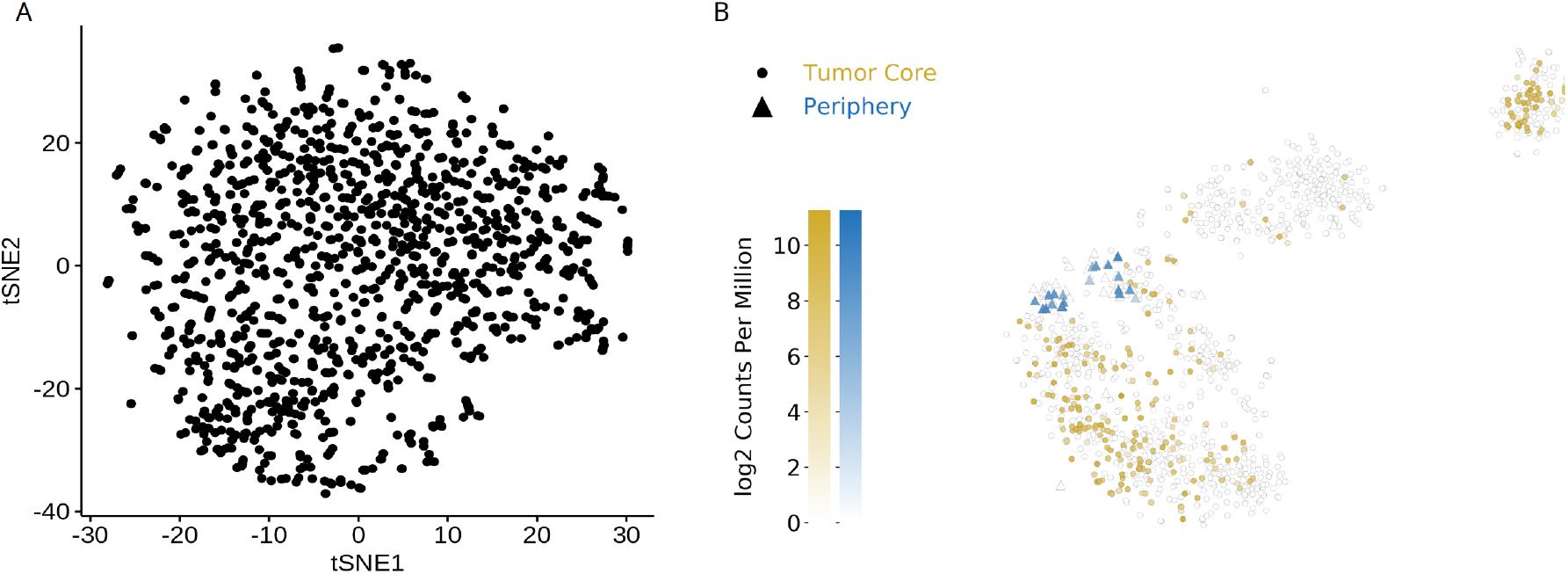
Clustering of IDH-A cells and Tlx expression pattern in human GBM cells. **A**, tSNE projection of MGH61 malignant cells (Andrew Venteicher, et al., 2017). Processed data were downloaded from GEO and highly variable genes were used for tSNE embedding. **B**, Tlx expression profile in human GBM (Spyros Darmanis, et al., 2017). Modified from http://www.gbmseq.org.

