## Supplementary Theory for "Exponential Growth of Glioblastoma *In Vivo* Driven by Rapidly Dividing and Outwardly Migrating Cancer Stem Cells"

August 2, 2019

#### Contents

**T1 Code availability** **2**

**T2 Phenomenological Modelling of Tumour Growth Curves** **2**

**T3 Three-dimensional Tumour Growth Simulation** **3**

**T4 Mathematical Model of Glioblastoma Growth** **7**

**T5 Describing Experiments with the Mathematical Model** **8**

#### T1 Code availability

All code discussed in this supplement including experimental and simulated data required for its execution is available at https://github.com/LiBuchauer/exponential_GBM.

All computations were performed using python 3.5. This work relies on the following external packages:

- numpy [van der Walt et al., 2011]
- matplotlib [Hunter, 2007]
- seaborn [Waskom et al., 2018]
- lmfit [Newville et al., 2014]
- pandas [McKinney, 2010]
- scipy [Jones et al., 2001]
- emcee [Foreman-Mackey et al., 2013]
- corner [Foreman-Mackey et al., 2016]
- Cython [Behnel et al., 2011]

#### T2 Phenomenological Modelling of Tumour Growth Curves

Figures T1 to T3 show the bioluminescence imaging (*BLI*) curves recorded from 23 glioblastoma-bearing animals over periods of several weeks. We are interested in the dynamics of this growth phase and want to make use of it for estimating parameters of glioblastoma growth in section T6. A wide range of mathematical models have been proposed for modelling tumour growth and commonly employed models include exponential, Gompertzian and power-law growth [Wallace and Guo, 2013, Talkington and Durrett, 2015, Murphy et al., 2016]. Here, we have fitted these three models to the 23 experimental time series in figures T1 to T3 using the Nelder-Mead method.

In the simplest case, the exponential model, growth is governed by rate $\lambda_0$,

$$N_{\mathrm{exp}}(t) = N_0 \mathrm{e}^{\lambda_0 t}, \tag{1}$$

where $N_{\mathrm{exp}}(t)$ describes the number of cells at time $t$ assuming $N_0$ cells at $t = 0$. The Gompertzian growth function is given by

$$N_{\mathrm{Gom}}(t) = K_0 \mathrm{e}^{\log(N_0/K_0)\exp(-g_0 t)}, \tag{2}$$

where $N_{\mathrm{Gom}}(t)$ is again the number of cells at time $t$ when having started from $N_0$ at $t = 0$. The Gompertzian curve describes a sigmoidal behaviour where growth slows down at the end of the process - this behaviour is mediated by the carrying capacity $K_0$. The parameter $g_0$ is related to the proliferation speed of cells. Lastly, the power-law model incorporates the idea that only the cells on the surface of a solid tumour are able to proliferate, for example because of nutrient constraints in the core. The tumour volume $V_{\mathrm{pow}}(t)$ at time $t$ thus depends on the initial radius of the tumour, $R_0$, the linear growth rate $r_0$ or the tumour's radius. This results in the function

$$V_{\text{pow}}(t) = \frac{4\pi}{3}\left(r_0 t + R_0\right)^3 . \tag{3}$$

While the power-law model yields significantly worse fit results than exponential and Gompertzian growth curves by measure of comparing the resulting $\chi^2$ values, these two models perform similarly well. However, the Gompertzian model requires an additional parameter as compared to the exponential growth model. Using Akaike's information criterion [Akaike, 1998], which includes a penalty for more complex models, we find that the exponential model is preferable in 18 out of the 23 cases analysed. In the remaining five cases, Gompertzian growth fit the data best. These results suggest that while glioblastoma growth reaches a capacity limit eventually, our study focuses on the phase of tumour expansion where growth is still exponential.

In order to describe the mean dynamics of this exponential growth phase, we take the mean of the 23 fitted values of $\lambda_0$ (shown in figure T1) resulting in

$$\lambda_0 \pm \sigma_{\lambda_0} = (0.21 \pm 0.02)\ \text{day}^{-1} ,$$

where $\sigma_{\lambda_0}$ is the associated standard error of mean.

When fitting each of the three models in equations (1), (2) and (3) to the data, the initial values $N_0$ (respectively $R_0$) were allowed to vary freely in $\mathbb{R}^+$ in addition to the model parameters of interest. This absorbs much of the animal-to-animal variability which may be due to different numbers of initially transformed malignant cells or the positions of the tumour in the brain with respect to the skull and measurement devices. It is likewise for this reason that their values carry no information accessible or relevant to the current discussion.

#### T3 Three-dimensional Tumour Growth Simulation

In order to explore the effects of different cellular migration behaviours on growth curves of solid tumours, we implemented a tumour growth simulation on a three-dimensional Cartesian grid. This simulation comprises a self-renewing cell type (type $A$) and a derived cell type which is unable to divide (type $B$) as well as behavioural rules associated with cell types $A$ and $B$.

Self-renewing cells of type $A$ can divide symmetrically ($A \longrightarrow 2A$, rate $\lambda_S$) or asymmetrically ($A \longrightarrow A + B$, rate $\lambda_A$) while cells of type $B$ are not proliferative. Neither cell type dies in this simplified scenario. In addition, according to the user's choice, one or both of the cell types can migrate around the grid. This cellular movement happens towards the edge of the tumour; the direction of the moves is set to the vector between the centre of mass of the tumour and the current position of the cell. Faster migration and thus dispersal into the surrounding area can be enforced by setting cells to take more than one step into the chosen direction. In the results displayed in the main text, this dispersal factor was set to 3. Uniform integer random noise in the range $[-2, 2]$ was added to each component of the migration vector independently in order to avoid restriction of movement to the 26 Moore axes. The event rates used in the main text are $\lambda_S = 0.23$ and $\lambda_A = 0.65$; all migration decisions happen with rate $\rho = 0.4$ irrespective of migration scenario.

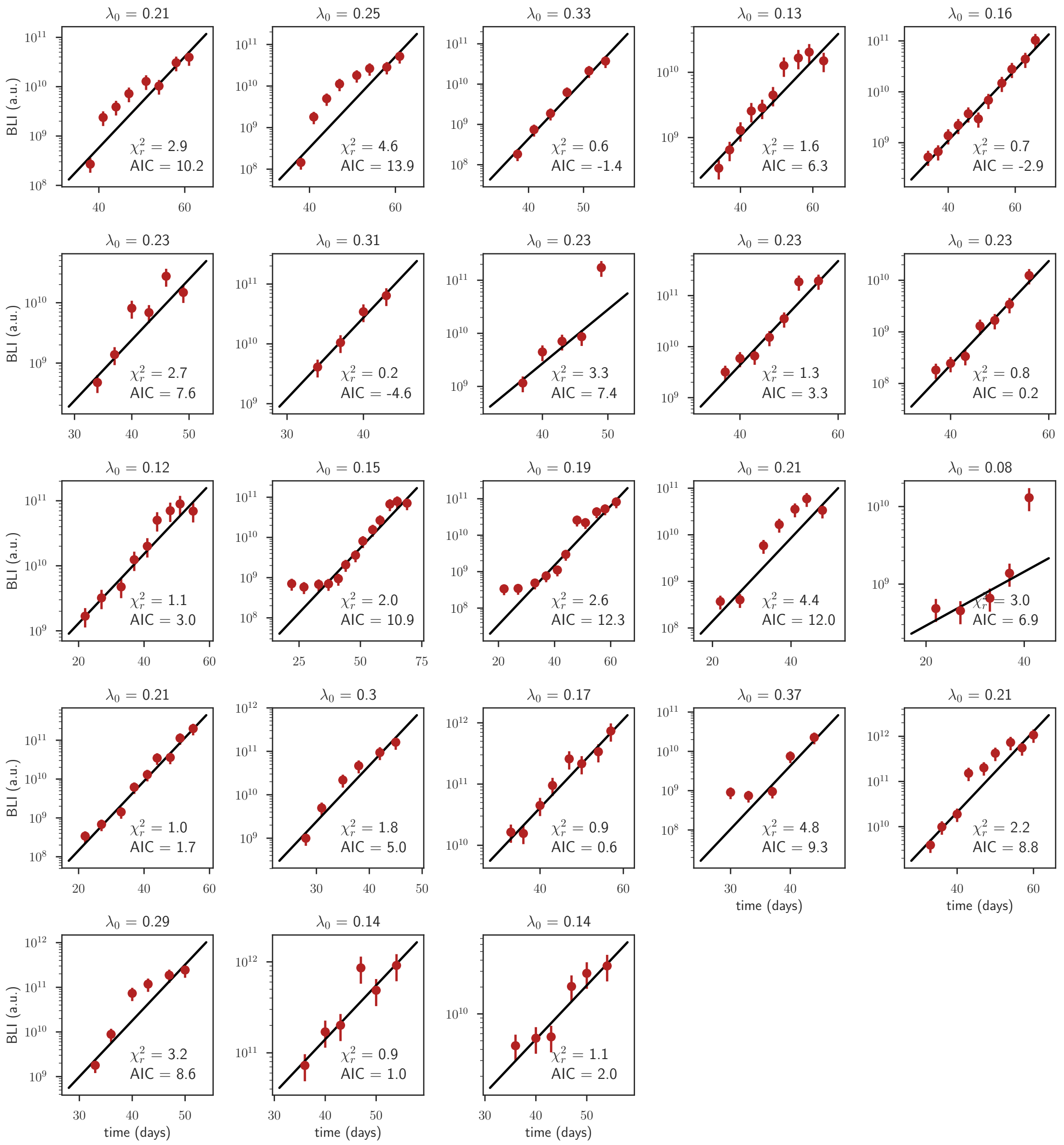

Figure T1: Time course of glioblastoma growth in 23 individual animals as shown by the increase of bioluminescence over time after birth (and induction of tumour growth) in days. Red dots show measured BLI values with an assumed relative error of $^1/_3$, black lines show best fits of the exponential model in equation (1). The resulting best fit values of $\lambda_0$ are given above each individual plot in units of day$^{-1}$.

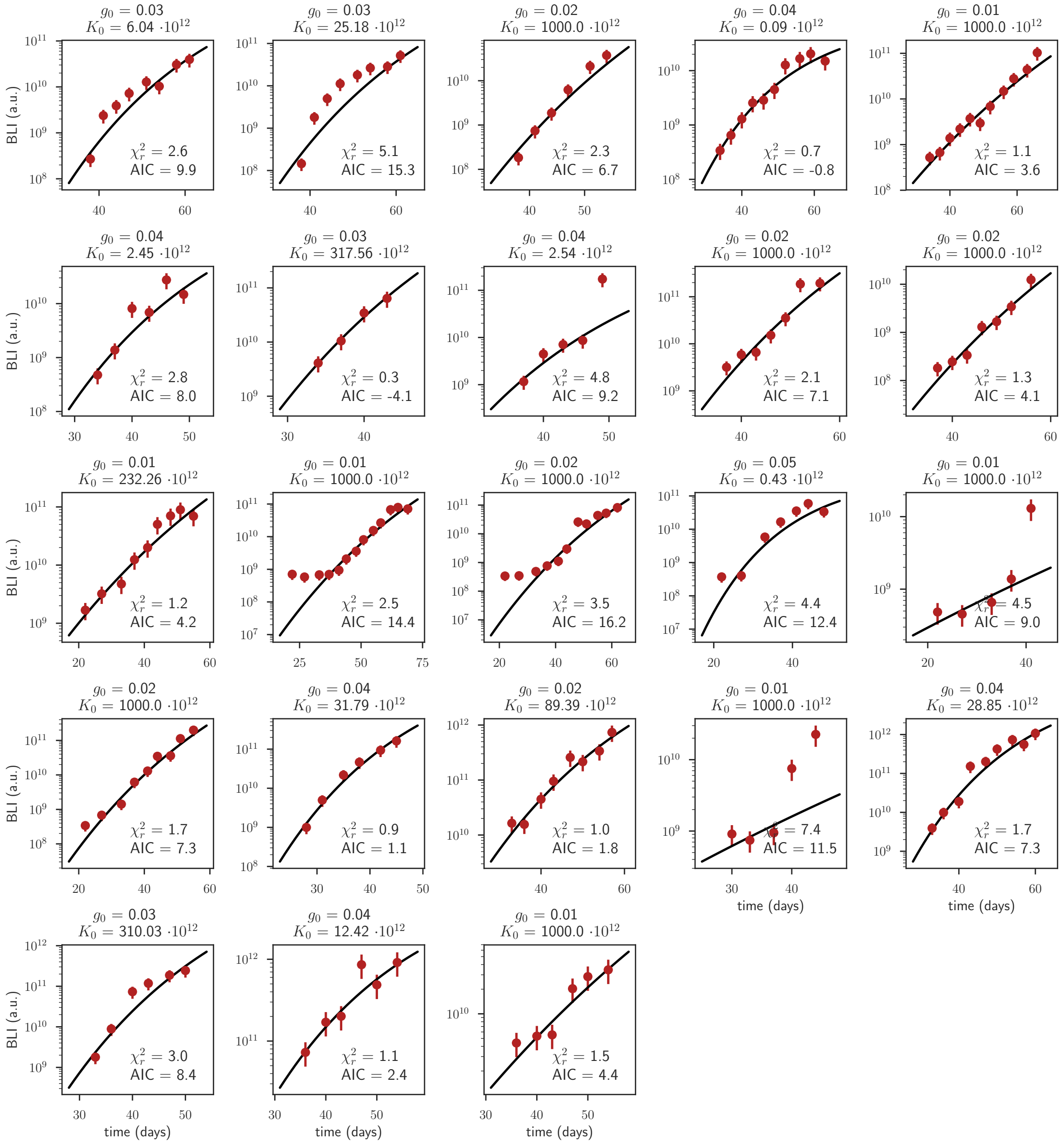

Figure T2: Time course of glioblastoma growth in 23 individual animals as shown by the increase of bioluminescence over time after birth (and induction of tumour growth) in days. Red dots show measured BLI values with an assumed relative error of $^1/_3$, black lines show best fits of the Gompertzian model in equation (2). The resulting best fit values of $K_0$ (without unit) and $g_0$ (units of day$^{-1}$) are given above each individual plot.

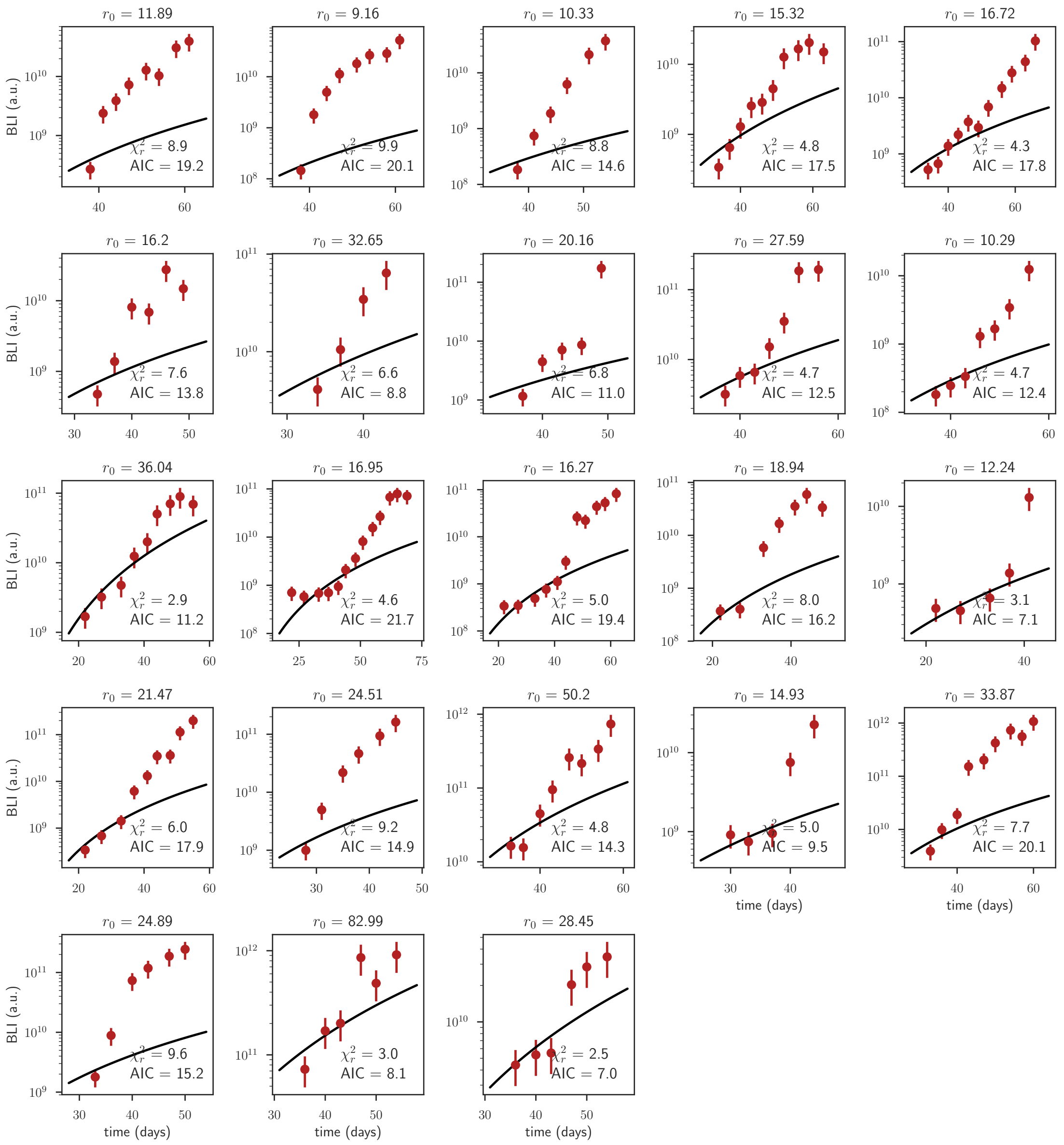

Figure T3: Time course of glioblastoma growth in 23 individual animals as shown by the increase of bioluminescence over time after birth (and induction of tumour growth) in days. Red dots show measured BLI values with an assumed relative error of $^1/_3$, black lines show best fits of the power-law model in equation (3). The resulting best fit values of $r_0$ are given above each individual plot in units of day$^{-1}$.

Algorithm 1 shows a compressed version of the simulation relying on Gillespie's stochastic simulation algorithm [Gillespie, 1977]. Building on simulation output, 3D images shown in the main text where rendered using a combination of mathematica for translating the occupancy grid produced by the python code into a 3D-image file and the open-source 3D computer graphics software blender.

**Algorithm 1** Simulation algorithm of three-dimensional tumour growth with migration.

**function** TUMOUR GROWTH(symmetric division rate, asymmetric division rate, migration rate(s), desired migration behaviour)
  Initialise an empty three-dimensional Cartesian grid.
  Put one cell of type $A$ into the middle of the grid.
  **while** $t_{\text{now}} < t_{\text{end}}$ **do**
    Based on a random number generator, the current cell population in the grid and the rates associated with the possible events, decide which type of event will happen next and after what time increment $\Delta t$.
    **if** event is symmetric division of cell type $A$ **then**
      Select one cell of type $A$ at random.
      **if** this cell has one or more empty neighbour sites **then**
        Choose a random empty neighbour and add a new cell of type $A$ there.
      **else**
        Do not divide.
    **else if** event is asymmetric division of cell type $A$ **then**
      Select one cell of type $A$ at random.
      **if** this cell has one or more empty neighbour sites **then**
        Choose a random empty neighbour and add a new cell of type $B$ there.
      **else**
        Do not divide.
    **else if** event is migration of cell type $A$ (or $B$) **then**
      Select one cell of type $A$ (or $B$) at random.
      Calculate how to get away from center of mass, add noise to the move, move the cell to its new position.
    Update time, $t_{\text{now}} \longleftarrow t_{\text{now}} + \Delta t$
  **return** the current occupancy of the grid with cells of type $A$ and $B$.

#### T4 Mathematical Model of Glioblastoma Growth

As introduced in the main text, we here choose to model glioblastoma growth with a simple mathematical model comprising three cell types: cancer stem cells ($S$), their actively dividing progeny ($P$) and terminally differentiated cells which are no longer able to proliferate ($D$) and dub it *SPD*-model. We include the following cellular processes into our mathematical analysis: symmetric division of stem cells ($S \longrightarrow 2S$) with rate $\lambda_S$, asymmetric division of stem cells ($S \longrightarrow S + P$) with rate $\lambda_A$, symmetric division of progeny ($P \longrightarrow 2P$) with rate $\lambda_1$, differentiation of an actively dividing progeny cell into a terminally differentiated cell ($P \longrightarrow D$) with rate $d_1$ and finally loss of terminally differentiated cells ($D \longrightarrow \emptyset$) with rate $\mu_1$. Assuming mass-action kinetics, the dynamics of this system are formalised by the following set of ordinary differential equations in the limit of large cell numbers,

$$\begin{aligned} \dot{S}(t) &= \lambda_S S(t) \\ \dot{P}(t) &= \lambda_A S(t) + (\lambda_1 - d_1)P(t) \\ \dot{D}(t) &= d_1 P(t) - \mu_1 D(t)\,. \end{aligned} \tag{4}$$

Explicit solutions for this system of equations are readily available. Assuming tumour growth to start from one stem cell at time point $t = 0$ (initial conditions $S(0) = 1, P(0) = 0, D(0) = 0$), we find

$$\begin{aligned} S(t) &= \mathrm{e}^{\lambda_S t} \\ P(t) &= \frac{\lambda_A}{\lambda_S + d_1 - \lambda_1}\left(\mathrm{e}^{\lambda_S t} - \mathrm{e}^{(\lambda_1 - d_1)t}\right) \\ D(t) &= \xi\left[(d_1 - \lambda_1 - \mu_1)\mathrm{e}^{\lambda_S t} + (\lambda_S + \mu_1)\mathrm{e}^{(\lambda_1 - d_1)t} + (\lambda_1 - d_1 - \lambda_S)\mathrm{e}^{-\mu_1 t}\right], \\ \xi &= \frac{\lambda_A d_1}{(\lambda_S + \mu_1)(\lambda_1 - d_1 + \mu_1)(\lambda_S + d_1 - \lambda_1)}\,. \end{aligned} \tag{5}$$

Because of the experimental observation by [Zhu et al., 2014] that $\mathrm{Tlx}^-$ cells (cells of type $P$ and $D$ in the language of the model) cannot in general sustain tumour growth by themselves, we require that cells of type $P$ always be lost via differentiation at least as fast as they divide,

$$\lambda_1 - d_1 \leq 0\,. \tag{6}$$

Revisiting equation (5) with equation (6) in mind, we see that the second term in $P(t)$ as well as the second and third term in $D(t)$ become irrelevant at later times because their exponents assume negative values for all times. The only remaining terms are governed by the symmetric division rate of stem cells $\lambda_S$ for all three populations. Defining the overall tumour dynamics $N(t)$ via

$$N(t) = S(t) + P(t) + D(t)\,, \tag{7}$$

we thus find the effective growth rate of the whole tumour to be governed only by the symmetric division rate of stem cells for $t \gg 0$,

$$N(t) \propto \mathrm{e}^{\lambda_S t}\,. \tag{8}$$

As a further property of this model, the fractional composition of the tumour bulk reaches a steady state dependent on the five rates defining the model even though the tumour cell population grows exponentially for $\lambda_S > 0$. These stationary fractions are given by

$$\begin{aligned} \langle\frac{S}{N}\rangle &= \kappa(\lambda_S + d_1 - \lambda_1)(\lambda_S + \mu_1) \\ \langle\frac{P}{N}\rangle &= \kappa\lambda_A(\lambda_S + \mu_1) \\ \langle\frac{D}{N}\rangle &= \kappa\lambda_A d_1\,, \\ \kappa &= \frac{1}{(\lambda_S + \lambda_A - \lambda_1)(\lambda_S + \mu_1) + d_1(\lambda_A + \lambda_S + \mu_1)}\,. \end{aligned} \tag{9}$$

#### T5 Describing Experiments with the Mathematical Model

Here we describe how several experimentally observed quantities can be simulated using the basic mathematical model described in the section T4. The experiments discussed in sections T5.1, T5.2 and T5.3 are used for parameter estimation in section T6.

##### T5.1 BLI Growth Curves

Since the phenomenological growth curve description of equation (1) matches the functional form of the mechanistically derived model growth curve of equation (8), in order to describe the BLI tumour growth data adequately with our model, we must seek to minimise the difference between $\lambda_S$ and the experimental value of $\lambda_0$ derived above. Assuming the experimental $\lambda_0$ to stem from a Gaussian distribution, this results in the following likelihood function for the BLI data given the model parameter $\lambda_S$:

$$\mathscr{L}_{BLI}(\lambda_0, \sigma_{\lambda_0}|\lambda_S) = \frac{1}{\sqrt{2\pi\sigma^2_{\lambda_0}}} \exp\left(-\frac{(\lambda_0 - \lambda_S)^2}{2\sigma^2_{\lambda_0}}\right). \tag{10}$$

Thus, we find for the logarithm of the likelihood

$$\log \mathscr{L}_{BLI}(\lambda_0, \sigma_{\lambda_0}|\lambda_S) = \text{const.} - \frac{(\lambda_0 - \lambda_S)^2}{2\sigma^2_{\lambda_0}}, \tag{11}$$

which will form part of the composite log-likelihood function at the basis of parameter estimation in section T6.

##### T5.2 Decline of Proliferation Index in the Progeny Compartment

In this experiment, the development of the proliferative capacity of $\text{Tlx}^-$ (progeny) cells was measured over time. To this end, viral vectors carrying the gene for the red fluorescent protein dsRed were administered approximately 30 days after birth, thus genetically labelling all currently dividing progeny cells. In the following, the time point of this labelling defines $t_{\text{dsRed}}$, the start of experiment and simulation. A total of 13 animals were then sacrificed at time points $t_{\text{dsRed}} + \Delta t_i$, $\Delta t_i \in [1 \text{ day} \ldots 13 \text{ days}]$, and tumour sections were stained for the proliferation marker Ki67 (main text figure 5D). As an error estimate, the sample standard deviation was calculated on the time point with the largest sample size available (day 6, $n = 3$) and used as an approximation to the population standard deviation $\sigma_{Ki67}$ of all time points. The SEM estimate for each time point $\Delta t_i$ was then calculated using the number of animals available at this time point, $n(\Delta t_i)$, giving $\sigma_{\mathscr{F}}(\Delta t_i) = \sigma_{Ki67}/\sqrt{n(\Delta t_i)}$.

To simulate this experiment, we first evolve the tumour model given by equations (4) until day 30, the approximate starting point of the real experiment. However, the exact starting point of the simulated experiment does not influence the outcome, as - like in the experimental case - we are interested only in the fraction of cells still proliferating after time intervals $\Delta t$. The time course of this fraction is independent of the starting time because all rates in the model are assumed to be stationary.

Let the viral vector introduce dsRed into all cells of type $P$ at $t_{\text{dsRed}} = 30$ days while not labelling any stem cells or terminally differentiated cells. This gives as initial conditions for our experiment:

$$\begin{aligned} S_{\text{dsRed}}(t_{\text{dsRed}}) &= 0 \\ P_{\text{dsRed}}(t_{\text{dsRed}}) &= P(30 \text{ days}) \\ D_{\text{dsRed}}(t_{\text{dsRed}}) &= 0\,. \end{aligned} \tag{12}$$

Using these initial conditions, we solve equations (4) numerically resulting in solutions $P_{\text{dsRed}}(t_{\text{dsRed}} + \Delta t)$ and $D_{\text{dsRed}}(t_{\text{dsRed}} + \Delta t)$. $S_{\text{dsRed}}(t_{\text{dsRed}} + \Delta t) = \text{const.} = 0$ because no stem cells incorporate dsRed and there is no dedifferentiation in the model. The modelled proliferative fraction $\mathscr{F}_{\text{mod}}^{\text{Ki67}}$ within the dsRed population at $\Delta t$ is then readily calculated as

$$\mathscr{F}_{\text{mod}}^{\text{Ki67}}(\Delta t) = \frac{P_{\text{dsRed}}(t_{\text{dsRed}} + \Delta t)}{P_{\text{dsRed}}(t_{\text{dsRed}} + \Delta t) + D_{\text{dsRed}}(t_{\text{dsRed}} + \Delta t)} \,. \tag{13}$$

In analogy to equations (10) and (11), we can now derive the parameter-dependent logarithmic likelihood of the experimental Ki67-fractions as shown in main text figure 5D for a given set of model parameters $\theta = \{\lambda_S, \lambda_A, \lambda_1, d_1, \mu_1\}$ to

$$\log \mathscr{L}_{\text{Ki67}}(\{\mathscr{F}_{\text{exp}}^{\text{Ki67}}, \sigma_{\mathscr{F}}\}|\theta) = \text{const.} - \sum_i \frac{\left(\mathscr{F}_{\text{exp}}^{\text{Ki67}}(\Delta t_i) - \mathscr{F}_{\text{mod}}^{\text{Ki67}}(\Delta t_i;\theta)\right)^2}{2\left(\sigma(\Delta t_i)\right)^2} \,, \tag{14}$$

where the sum runs over all time points where experimental information is available.

##### T5.3 Tumour Composition Information

As discussed in section T4, the fractional decomposition of the tumour into its three populations $S$, $P$ and $D$ is stationary for $t \gg 0$ and is given by equation (9). The fraction of stem cells within the tumour bulk was assessed by flow cytometry using a mouse model which expresses RFP in all glioblastoma cells and additionally GFP in $\text{Tlx}^+$ cells (stem cells) while the fractions of proliferating and terminal progenitors are currently not accessible experimentally. Additionally, the fraction of actively proliferating cells within the tumour bulk ($\mathscr{F}_{\text{exp}}^{\text{prolif/bulk}}$) and the fraction of proliferating stem cells within the population of proliferating cells ($\mathscr{F}_{\text{exp}}^{\text{stem/prolif}}$) were measured by a combination of staining for the proliferation marker Ki67 and the mouse model co-expressing GFP and Tlx. The value of $\mathscr{F}_{\text{exp}}^{\text{stem/prolif}}$ was previously published in [Zhu, 2013] as the fraction of tumour cells stained positively for the proliferation marker PCNA; the value for $\mathscr{F}_{\text{exp}}^{\text{prolif/bulk}}$ shown here is a weighted average of values from [Zhu, 2013, Zhu et al., 2014] and new data reported here, all based on Ki67-stainings:

$$\begin{aligned}
\mathscr{F}_{\text{exp}}^{\text{stem/bulk}} &= (0.23 \pm 0.04)\% \,, \\
\mathscr{F}_{\text{exp}}^{\text{prolif/bulk}} &= (0.22 \pm 0.04)\% \,, \\
\mathscr{F}_{\text{exp}}^{\text{stem/prolif}} &= (0.09 \pm 0.02)\% \,.
\end{aligned}$$

These experimental ratios and their model counterparts are visualised in main text figure 5C.

In order to construct equivalent ratios from our mathematical model, we must make assumptions about the proliferative action of the three populations $S$, $P$ and $D$. Fully differentiated cells of type $D$ are not proliferative per model definition. Similarly, all proliferating progeny of type $P$ is actively dividing by construction of the model. The case of stem cells $S$ however is less clear, as we know stem cells both in the neural case and in general to enter and exit the cell cycle depending on external and internal cues, thus switching between active and quiescent state [Li and Clevers, 2010, Ponti et al., 2013, Bond et al., 2015]. Here, we have chosen not to include two separate stem cell populations into the model because we have insufficient information for informing all rates resulting from an additional cell population. Nevertheless, cell cycle analysis in our tumour model indicates that $f_{\text{G}_0} = (83 \pm 6)\%$ of $\text{Tlx}^+$ cells are in the $\text{G}_0$-phase at any one time [Zhu et al., 2014]. We make use of this information directly in modelling the two experimentally observed ratios introduced above without an explicit quiescent population by assuming the remaining $(1 - f_{G_0})$ to be actively proliferating. As a result, we define for comparison with experimental observations

$$\begin{aligned}
\mathscr{F}_{\text{mod}}^{\text{stem/bulk}} &= \lim_{t\to\infty} \frac{S(t)}{N(t)}\,, \\
\mathscr{F}_{\text{mod}}^{\text{prolif/bulk}} &= \lim_{t\to\infty} \frac{(1-f_{G_0})S(t)+P(t)}{N(t)}\,, \\
\mathscr{F}_{\text{mod}}^{\text{stem/prolif}} &= \lim_{t\to\infty} \frac{(1-f_{G_0})S(t)}{(1-f_{G_0})S(t)+P(t)}\,.
\end{aligned} \tag{15}$$

The stationarity of the second and third model ratios follows directly from the stationarity of the subpopulation fractions given in equation (9). Their values were calculated numerically.

As in the previous sections, these considerations lead to a parameter-dependent logarithmic likelihood contribution from the tumour composition data of the form

$$\begin{aligned}
\log \mathscr{L}_{\text{comp}}(\mathscr{F}_{\text{exp}}^{\text{s/b}}, \mathscr{F}_{\text{exp}}^{\text{p/b}}, \mathscr{F}_{\text{exp}}^{\text{s/p}}, \sigma_{\mathscr{F}_{\text{s/b}}}, \sigma_{\mathscr{F}_{\text{p/b}}}, \sigma_{\mathscr{F}_{\text{s/p}}}|\theta) = \text{const.} - \frac{(\mathscr{F}_{\text{exp}}^{\text{s/b}} - \mathscr{F}_{\text{mod}}^{\text{s/b}}(\theta))^2}{2\sigma^2_{\mathscr{F}_{\text{s/b}}}} \\
- \frac{(\mathscr{F}_{\text{exp}}^{\text{p/b}} - \mathscr{F}_{\text{mod}}^{\text{p/b}}(\theta))^2}{2\sigma^2_{\mathscr{F}_{\text{p/b}}}} - \frac{(\mathscr{F}_{\text{exp}}^{\text{s/p}} - \mathscr{F}_{\text{mod}}^{\text{s/p}}(\theta))^2}{2\sigma^2_{\mathscr{F}_{\text{s/p}}}}\,,
\end{aligned} \tag{16}$$

which is used for joint parameter estimation in section T6.

### T6 Parameter Estimation: Method and Results

In order to calibrate the model introduced in section T4 to the data using the likelihoods derived in section T5, we use a Bayesian framework in which we approximate the posterior distribution of model parameters with Markov chain Monte Carlo (MCMC) sampling. This section comprises a short summary of the principles of Bayesian parameter estimation (section T6.1), a summary of how MCMC sampling was employed (section T6.2) and an overview of the resulting parameter estimates and their confidence bounds (section T6.3).

#### T6.1 Bayesian Parameter Estimation

At the heart of Bayesian parameter estimation (see for example [Sorensen and Gianola, 2002, MacKay, 2005, Box and Tiao, 2011]) lies Bayes' Theorem which provides the probability distribution of parameters $\theta$ as a function of the available experimental data $D$, a chosen model topology $\mathscr{H}$, an error model and distribution conveying prior information on the distribution of parameters,

$$P(\theta|D,\mathscr{H}) = \frac{\mathscr{L}(D|\theta,\mathscr{H})\,\pi(\theta|\mathscr{H})}{P(D|\mathscr{H})}\,. \tag{17}$$

This resulting probability distribution $P(\theta|D,\mathscr{H})$ over parameters is called *posterior distribution* or simply *posterior*. Data $D$ as well as model and error assumptions enter the calculation via the likelihood function $\mathscr{L}(D|\theta,\mathscr{H})$, potential prior information is encoded in the *prior distribution* (or *prior*) $\pi(\theta)$. The normalising constant $P(D|\mathscr{H})$ naturally gives the total probability of the data given the chosen model when taking into account all possible parameter sets $\theta$,

$$P(D|\mathscr{H}) = \int \mathrm{d}\theta\, \mathscr{L}(D|\theta,\mathscr{H})\,\pi(\theta|\mathscr{H})\,. \tag{18}$$

We have previously introduced all components necessary for making use of Bayes' theorem as shown in equation (17) - a mathematical model in section T4 and experimental data and associated error models in section T5. Specifically we have given likelihood functions for all datasets in equations (11), (14) and (16). Since these experiments are independent of each other, we arrive at a combined likelihood function $\mathscr{L}_{\text{total}}$ by multiplying the individual likelihood contributions. This gives, up to a constant, a total logarithmic likelihood of

$$\begin{aligned}\log \mathscr{L}_{\text{total}}(D|\theta) = \; & \log \mathscr{L}_{BLI}(\lambda_0, \sigma_{\lambda_0}|\lambda_S) + \log \mathscr{L}_{\text{Ki67}}(\{\mathscr{F}^{\text{Ki67}}_{\text{exp}}, \sigma_{\mathscr{F}}\}|\theta) + \\ & \log \mathscr{L}_{\text{comp}}(\mathscr{F}^{\text{s/b}}_{\text{exp}}, \mathscr{F}^{\text{p/b}}_{\text{exp}}, \mathscr{F}^{\text{s/p}}_{\text{exp}}, \sigma_{\mathscr{F}_{\text{s/b}}}, \sigma_{\mathscr{F}_{\text{p/b}}}, \sigma_{\mathscr{F}_{\text{s/p}}}|\theta),\end{aligned} \tag{19}$$

where $\theta = \{\lambda_S, \lambda_A, \lambda_1, d_1, \mu_1\}$ as previously. As for the prior distribution over these parameters, we assume them to be uniformly distributed within biologically reasonable bounds. Furthermore we assume prior independence of parameters allowing us to give distributions $\pi^{\dagger}(\beta)$ on each parameter separately. For convenience, we choose the same bounds for all five rates (0 and 4 day$^{-1}$),

$$\pi^{\dagger}(\beta) = \begin{cases} 0 & \text{for } \alpha < 0, \\ \frac{1}{4}, & \text{for } 0 \leq \alpha \leq 4 \\ 0 & \text{for } \alpha > 4 \end{cases}, \quad \text{for } \beta \text{ in } \{\lambda_S, \lambda_A, \lambda_1, d_1, \mu_1\} \tag{20}$$

and construct $\pi(\theta)$ as the product of the five individual priors. As discussed above (equation (6)), we require $d1 \leq \lambda_1$ because of previous observations and thus assign a prior probability of zero to any parameter set that does not fulfil this constraint. As we will see below, our data yield fully identifiable parameter estimates far away from the somewhat arbitrarily chosen upper bound of 4 /day, thus confirming its minor importance for our inference. After dropping all constants from equation (17) in our context, we redefine

$$P(\theta|D) = \mathscr{L}(D|\theta)_{\text{total}}\, \pi^{\dagger}(\lambda_S)\pi^{\dagger}(\lambda_A)\pi^{\dagger}(\lambda_1)\pi^{\dagger}(d_1)\pi^{\dagger}(\mu_1) \tag{21}$$

for the following discussion.

##### T6.2 MCMC Sampling of the Posterior Distribution

Because of the complicated composite likelihood function in equation (19), parts of which are evaluated numerically, we approximate the posterior parameter distribution $P(\theta|D)$ using an MCMC-approach. In general, MCMC-methods construct a Markov chain whose equilibrium distribution is the requested probability distribution [Sorensen and Gianola, 2002]. Samples taken from this chain after it has reached equilibrium may thus be regarded as samples of the target distribution. The calculations underlying this work have been performed in the programming language python and rely on the MCMC-package emcee [Foreman-Mackey et al., 2013]. This package uses an affine-invariant ensemble sampling method [Goodman and Weare, 2010], which runs many Markov chains in parallel and constructs informed choices for the next step of each single chain on the basis of the current positions of the others.

The result of MCMC-sampling the posterior parameter distribution defined by equations (19) and (20) is a 5-dimensional cloud of points encoding the probability associated with each region of parameter-space. Figure T4 shows all five 1-dimensional and all ten 2-dimensional marginal posterior distributions resulting for the five model parameters, i.e. (using parameters $\theta_1$ and $\theta_2$ as examples)

$$P(\theta_1|D) = \int \mathrm{d}\theta_2\, \mathrm{d}\theta_3\, \mathrm{d}\theta_4\, \mathrm{d}\theta_5\, P(\theta|D) \tag{22}$$

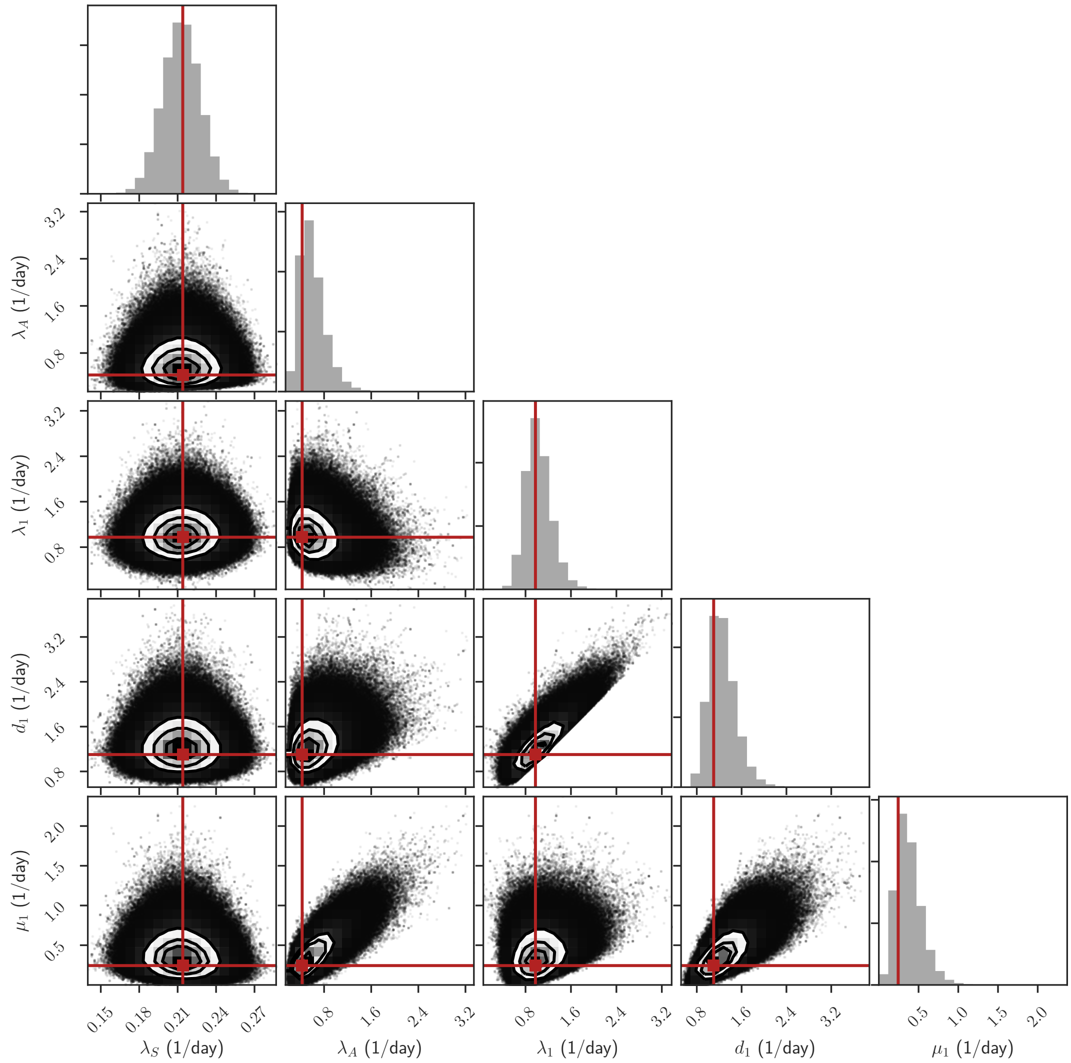

Figure T4: One- and two-dimensional projections of the posterior probability distribution of the model parameters $\lambda_S$, $\lambda_A$, $\lambda_1$, $d_1$ and $\mu_1$. The red lines indicate the position of the components of the most probable parameter set $\theta_{\mathrm{MAP}}$. The components of the joint modal vector do not generally coincide with the marginal modes as is obvious for $\lambda_A$ and $\mu_1$ here in particular.

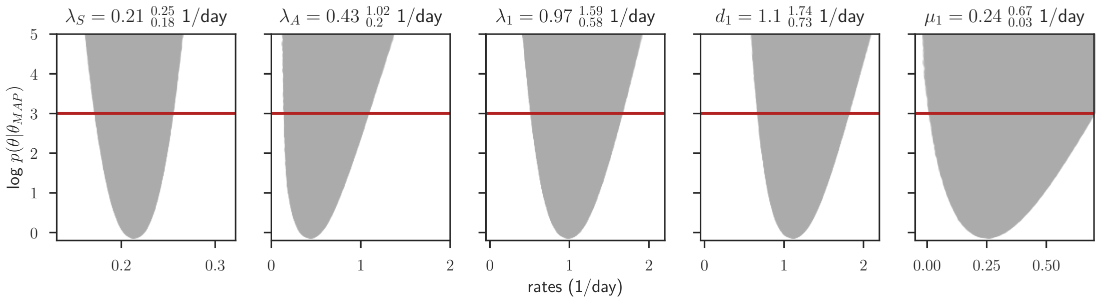

Figure T5: Posterior profiles of the five model parameters $\lambda_S, \lambda_A, \lambda_1, d_1$ and $\mu_1$ as discussed in section T6.3. The logarithm of the relative probability as defined in equation (25) is shown on the $y$-axis, with values above the red line corresponding to less than 5% of the probability associated with the best parameter set. The shaded area shows which relative probabilities are accessible for a given parameter value (shown on the $x$-axis). We define the parameter values where the red line cuts the shaded area as boundaries of the credible region of 5% relative probability and give the associated values in table T1.

for the marginal posterior of parameter $\theta_1$ and

$$P(\theta_1, \theta_2|D) = \int \mathrm{d}\theta_3\, \mathrm{d}\theta_4\, \mathrm{d}\theta_5 P(\theta|D)\,. \tag{23}$$

It is important to note that, in general, the components of the joint modal vector (mode of the full $n$-dimensional posterior, $\theta_{\mathrm{MAP}}$), are not equal to the modes of the $n$ marginal distributions [Sorensen and Gianola, 2002]. This can also be observed in figure T4, where the red lines have been added to indicate the components of the full modal vector - deviations from the marginal modes in one dimension are evident especially for $\lambda_A$ and $\mu_1$.

#### T6.3 MAP Parameter Estimates and Confidence Intervals

In the main text, we quote the maximum *a posteriori* (MAP) parameter set, i.e. the mode of the posterior, as a point estimate summarising the full parameter distribution shown in figure T5. This choice of estimator is linked to the decision-theoretic loss function $G_x(x_0; x) = \delta(x_0 - x)$, where $\delta$ is Dirac's delta function [Jaynes et al., 2003]. With this loss function,

$$\theta_{\mathrm{MAP}} = \arg\max_{\theta} \int \delta(\theta - \theta')P(\theta'|D)\mathrm{d}\theta'. \tag{24}$$

In practice, we make use of the probability values associated with each sampled set of the MCMC-chain for finding the most probable set $\theta_{\mathrm{MAP}}$ and defining credible regions around it. As this method is obviously vulnerable to random effects governing the sampling procedure, we have simulated MCMC-chains long enough to ascertain two stable decimal places for all values in $\theta_{\mathrm{MAP}}$ when repeating the entire approximation of the posterior. The results for the individual parameters are shown in figure T5 and can be found in table T1 as well.

In order to construct credible regions on these MAP estimates, we define the relative probability of every other parameter set $\theta$ as

$$p(\theta|\theta_{\mathrm{MAP}}) = \frac{\int \delta(\theta - \theta')P(\theta'|D)\mathrm{d}\theta'}{\int \delta(\theta_{\mathrm{MAP}} - \theta')P(\theta'|D)\mathrm{d}\theta'}\,. \tag{25}$$

| parameter | MAP estimate | $p(\theta\|\theta_{\mathrm{MAP}}) = 5\%$ credible region |
|---|---|---|
| $\lambda_S$ | 0.21 day$^{-1}$ | [0.18, 0.25] day$^{-1}$ |
| $\lambda_A$ | 0.43 day$^{-1}$ | [0.20, 1.02] day$^{-1}$ |
| $\lambda_1$ | 0.97 day$^{-1}$ | [0.58, 1.59] day$^{-1}$ |
| $d_1$ | 1.10 day$^{-1}$ | [0.73, 1.74] day$^{-1}$ |
| $\mu_1$ | 0.24 day$^{-1}$ | [0.03, 0.67] day$^{-1}$ |

Table T1: MAP parameter estimates and information regarding their credibility, for detailed discussion see text (sections T6.2, T6.3). For a graphical explanation of how credible regions were constructed see figure T5.

Using this definition, we can set a relative probability level $\alpha$ and regard as acceptable only those parameters within a credible region (CR)

$$\mathrm{CR} = \{\theta \mid p(\theta|\theta_{\mathrm{MAP}}) \geq \alpha\} \, . \tag{26}$$

We choose $\alpha = 0.05$, resulting in a cut-off of $|\log \alpha| = 3.00$ in logarithmic probability space. Figure T5 shows the application of this principle to our posterior distribution. For each of the five parameters, the figure indicates which relative probabilities can be assumed as a function of the chosen parameter if the other four parameters are allowed to vary freely. Thus, the realisations on the edge of the shaded regions in figure T5 represent the optimal accessible parameter set as function of each parameter in turn. The red line indicates where $\log p(\theta|\theta_{\mathrm{MAP}}) = 0.05$ - corresponding parameter bounds are defined where this line hits the accessible region of relative probability. Here we have used our MCMC-sampled posterior to determine these profiles which are hence only close approximations of the true optimal probabilities as a function of the fixed parameter. The credible regions resulting from this method are quoted in figure T5 and table T1.

In order to derive the confidence bounds on our simulations (projections of the MAP-parameter set into experimental space), we use only parameter sets from within the $p(\theta|\theta_{\mathrm{MAP}}) = 0.05$ - hypersurface. Several thousands of these parameter combinations are drawn and the experiments described in sections T5.1 to T5.3 are simulated with each of them. The resulting spread in simulation outcomes defines the bounds shown in main text figures 5C and D, which thus comprise all results accessible with parameter which are at least 1/20 as likely as the maximum probability estimate.

#### T7 Expanding the Model to Single Cell-Derived Clones

As described in the main text, section "Lineage Tracing Reveals a BTSC-Based Differentiation Hierarchy", individual cancer stem cells within a developed tumour can be genetically altered *in vivo* such that they start expressing one of four fluorescent proteins at random. The fluorescent protein is likewise expressed in all potential progeny of the initially labelled cell, thus enabling clonal tracing: when brain tumour slices are analysed by fluorescence microscopy a given time after the initial drug-induced labelling event, cells of the same fluorescent colour within close proximity of each other are likely derived from the same ancestor. Here, a total of eight animals were sacrificed at $\Delta t = \{5, 5, 10, 10, 20, 26, 37\}$ days after label induction and their fluorescent clones were quantified. From these datasets, we have derived the following summary statistics per animal (see main text figure 5E): mean number of cells per clone, coefficient of variation (CV) of the clone size and the relative fractions occupied by one-cell-clones and two-cell-clones within all quantified clones. Error bars accompanying data points of individual animals in this figure show estimates of the standard error derived using bootstrapping [Efron and Tibshirani, 1994, Boos, 2003].

We initially assumed the observed clonal growth dynamics to be governed by the same growth laws introduced and quantified for the population data in section T4. However, a comparison of the experimentally observed mean clone sizes with the model growth curve resulting from equations (5) and (7) using the MAP-parameter set $\theta_{\mathrm{MAP}}$ reveals a discrepancy that is both quantitative and qualitative: While the experimental mean clone size seems to reach a steady state by day 20 at the latest and never grows much beyond three cells per clone, the growth of the *SPD*-model starting from a single stem cell is exponential (see figure T6a). In order to resolve this contradiction, we refine the population model by allowing migration of cell types $S$ (stem cells) and $P$ (proliferating progeny) in a stochastic model extension. We define a migration event as a move of a cell far enough away from its clone of origin so that it is no longer identifiable as a clonal member by the observer but may instead be counted as an additional clone of its own. Figure T6b illustrates this principle and shows how it leads to more, but smaller, observed clones. We assign a new parameter, the migration rate $\beta$, to migration events. Stem cells and actively dividing progeny cells are assumed to leave their parent clone at equal speeds. Importantly, the original model retains validity for population-level processes, because cellular migration affects system properties only at the level of individual clones while leaving bulk properties like the total cell number or the cellular composition of the tumour unchanged.

For integrating migration at a single-cell level into the model, we again make use of stochastic simulation [Gillespie, 1977]. In order to include the emergence of new clones via migration into this framework, every time a cell of type $S$ or $P$ leaves the simulated clone, the size of this clone is reduced by one and the exact time of migration is recorded. After the original clone has been simulated until the required time, all clones spawned by migration and their offspring are likewise simulated from their respective time points of emergence. A Cython module of this simulation was written to enable efficient use in MCMC sampling. For the quantitative comparison between model and experiment, we focus on the four animals killed at later time points (days 20, 20, 26, 37). Because the mean clone size as well as the corresponding coefficient of variation and the fractions of size 1 and 2 clones seem to have reached saturation by day 20, we group these four animals into one steady state data set and calculate the mean of each of them for further comparison to the model. A conservative error estimate associated with these mean statistics is derived by Gaussian propagation of the four bootstrapped standard errors resulting in

$$
\begin{aligned}
\mu_{\mathrm{exp}} &= (2.8 \pm 0.5)\\
\mathrm{CV}_{\mathrm{exp}} &= (2.1 \pm 0.5)\\
\mathcal{F}_{\mathrm{exp}}^{\mathrm{size}\ 1} &= (0.56 \pm 0.04)\\
\mathcal{F}_{\mathrm{exp}}^{\mathrm{size}\ 2} &= (0.18 \pm 0.03)\,.
\end{aligned}
$$

To use this information in parameter estimation, the stochastic simulation starting from a single labelled cell has to be repeated enough times so that the true clone size distribution at the chosen day is sufficiently approached. Here, we find that when starting from 100 labelled cells and simulating until day 20, the four chosen summary statistics fall within a narrow in-

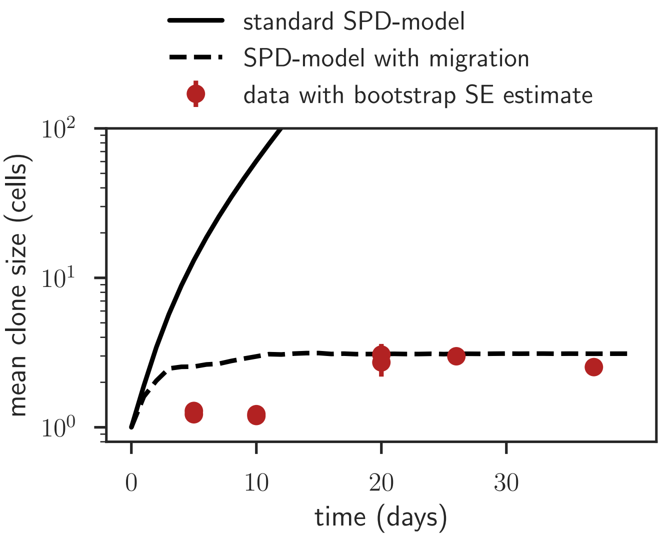

(a) Experimental mean clone size as a function of time (red dots) is not compatible with the bulk *SPD*-model from section T4, but can be explained when assuming that cellular migration leads to the observation of a higher number of smaller clones (see subfigure b).

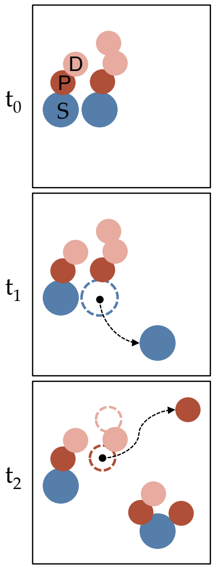

(b) Cellular migration leads to the observation of more and smaller clones.

Figure T6: Integrating migration processes into the model in order to resolve discrepant population and clonal growth patterns.

terval of less than 5% deviation from the mean upon multiple repetitions of the entire process (less than 1% for the fractions of size 1 and 2 clones). Additionally, a steady behaviour of the four statistics was reached by day 20 for all tested parameter combinations $\theta$ within the bounds given in table T1 supplemented by different migration rates $\beta$. Thus, because the simulation uncertainty of the steady state statistics is at least one order of magnitude below the experimental error, we continue to use the parameter estimation procedure introduced in section T6 as an approximation for this noisy optimisation problem - specifically, we define the likelihood of the clonal tracing data set in the same way as for the bulk datasets using the experimental error to weight the squared distance between model and experimental values. Uncertainties in the model values due to the stochastic nature of their simulation are omitted. In the following, the set of six parameters comprising $\lambda_S, \lambda_A, \lambda_1, d_1, \mu_1$ and $\beta$ is referred to as $\theta^\dagger$. We find for the approximated logarithmic likelihood contribution of the single-cell data:

$$\log \mathscr{L}_{\text{clones}}(\mu_{\text{exp}}, \text{CV}_{\text{exp}}, \mathscr{F}_{\text{exp}}^{\text{size }1}, \mathscr{F}_{\text{exp}}^{\text{size }2}, \sigma_\mu, \sigma_{\text{CV}}, \sigma_{\mathscr{F}^1}, \sigma_{\mathscr{F}^2} | \theta^\dagger) = \text{const.} - \frac{(\mu_{\text{exp}} - \mu_{\text{mod}}(\theta^\dagger))^2}{2\sigma_\mu^2} - \frac{(\text{CV}_{\text{exp}} - \text{CV}_{\text{mod}}(\theta^\dagger))^2}{2\sigma_{\text{CV}}^2} - \frac{(\mathscr{F}_{\text{exp}}^{\text{size }1} - \mathscr{F}_{\text{mod}}^{\text{size }1}(\theta^\dagger))^2}{2\sigma_{\mathscr{F}^1}^2} - \frac{(\mathscr{F}_{\text{exp}}^{\text{size }2} - \mathscr{F}_{\text{mod}}^{\text{size }2}(\theta^\dagger))^2}{2\sigma_{\mathscr{F}^2}^2}. \tag{27}$$

We add this logarithmic likelihood contribution to equation (19) to arrive at a new total likelihood including the single-cell data,

$$
\begin{aligned}
\log \mathscr{L}_{\text{total}}(D^\star|\theta^\dagger) = \; & \log \mathscr{L}_{BLI}(\lambda_0, \sigma_{\lambda_0}|\lambda_S) + \log \mathscr{L}_{\text{Ki67}}(\{\mathscr{F}^{\text{Ki67}}_{\text{exp}}, \sigma_{\mathscr{F}}\}|\theta) + \\
& \log \mathscr{L}_{\text{comp}}(\mathscr{F}^{\text{p/b}}_{\text{exp}}, \mathscr{F}^{\text{s/p}}_{\text{exp}}, \sigma_{\mathscr{F}_{\text{p/b}}}, \sigma_{\mathscr{F}_{\text{s/p}}}|\theta) + \\
& \log \mathscr{L}_{\text{clones}}(\mu_{\text{exp}}, \text{CV}_{\text{exp}}, \mathscr{F}^{\text{size }1}_{\text{exp}}, \mathscr{F}^{\text{size }2}_{\text{exp}}, \sigma_\mu, \sigma_{\text{CV}}, \sigma_{\mathscr{F}^1}, \sigma_{\mathscr{F}^2}|\theta^\dagger) \,.
\end{aligned}
\tag{28}
$$

We assume the prior distribution of the migration rate $\beta$ to match those of the other five parameters, given in equation (20) - a uniform distribution between 0 and 4/day. Performing the MCMC-sampling procedure in the same way as described in section T6 yields an approximation of the posterior distribution. Figure T7 shows the posterior profiles for all six parameters after inclusion of the single-cell tracing data. The previously determined values $\lambda_S, \lambda_A, \lambda_1, d_1$ and $\mu_1$ are in very good agreement with the earlier estimate, no significant shifts of the MAP values are observed. The newly introduced migration parameter $\beta$ is fully identifiable with a MAP value of 0.33 per day - this suggest that it takes cells of type $S$ and $P$ on average roughly 3 days to migrate far enough away that they cannot be identified as a member of the clone of origin any more by the observer.

### T8 Prediction of treatment outcomes

In this part, we discuss how predictions of the tumour volume's responses to two different forms of treatment were generated. Section T8.1 treats the response to chemotherapy, section T8.2 the response to knock-down of the stem cell factor Tlx.

#### T8.1 Chemotherapy with Temozolomide

Temozolomide (TMZ) is a chemotherapeutic anti-cancer drug that acts via DNA-methylation of mostly *N7* or *O6* guanine residues. During DNA replication this ultimately results in $G_2$/M cell cycle arrest and apoptosis [Zhang et al., 2012]. Thus, in the presence of TMZ, division attempts of cancerous cells result in cell death, while resting cells are not affected in the same way [Beier et al., 2008]. In order to model the effects of this drug, we alter the basic *SPD*-model given in equation (4) such as to incorporate them. Specifically, in the presence of the drug, we do not allow cell division to result in two daughter cells any more, but set it to lead to cell death instead. This concerns both symmetric and asymmetric division attempts of stem cells as well as divisions of progeny cells. The behaviour of fully differentiated cells in turn is not changed. This results in the following set of differential equations:

$$
\begin{aligned}
\dot{S}(t) &= -(\lambda_S + \lambda_A)\, S(t) \\
\dot{P}(t) &= -\lambda_1\, P(t) - d_1 P(t) \\
\dot{D}(t) &= d_1 P(t) - \mu_1 D(t)\,.
\end{aligned}
\tag{29}
$$

Of note, we have not introduced any additional parameters into the model here.

To mathematically replicate the treatment schedule, the tumour is first evolved in an undisturbed fashion until day 55, the approximate time point of experimental intervention. For this first phase, equations (4) are used. As treatment is experimentally applied for 10 days, equations (29) are used between day 55 and day 65 before returning to the undisturbed system thereafter. The model prediction resulting from this schedule is shown in main text figure 6C for the maximum MAP parameter set $\theta^\dagger_{\text{MAP}}$ and corresponding $\alpha = 0.05$ bounds. The bounds

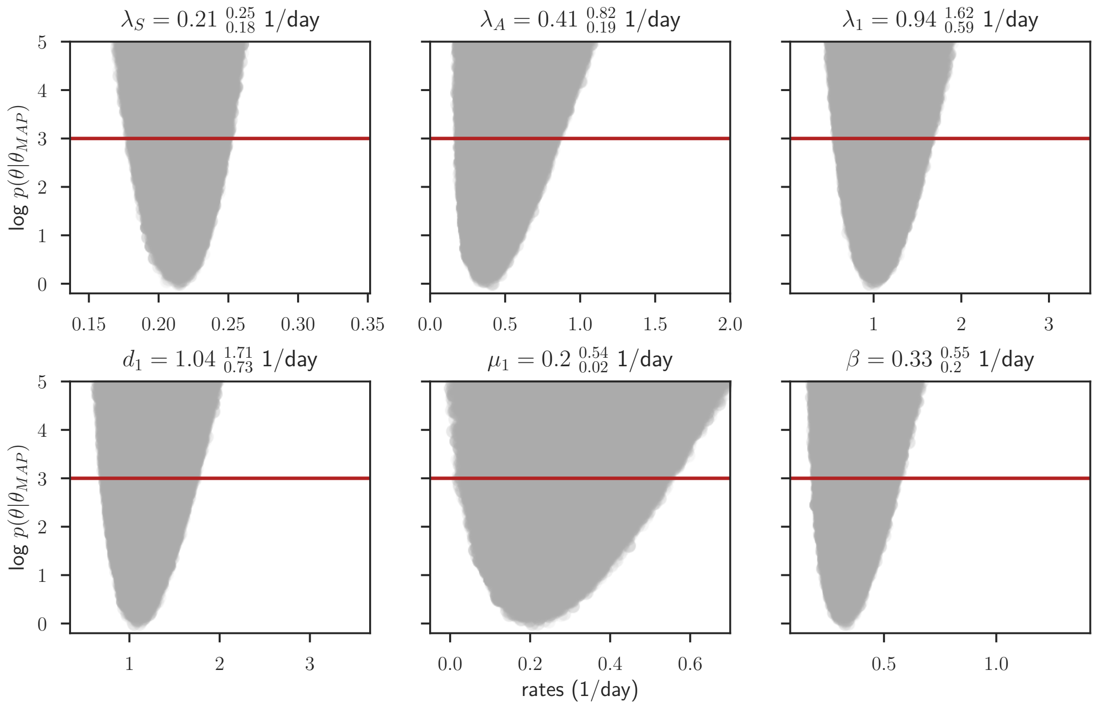

Figure T7: Posterior profiles of the six model parameters $\lambda_S, \lambda_A, \lambda_1, d_1, \mu_1, \theta$ and $\beta$ as discussed in section T7. The logarithm of the relative probability as defined in equation (25) is shown on the $y$-axis, with values above the red line corresponding to less than 5% of the probability associated with the best parameter set. As above (figure T5), the shaded area shows which relative probabilities are accessible for a given parameter value (shown on the $x$-axis). We define the parameter values where the red line cuts the shaded area as boundaries of the credible region of 5% relative probability. The fuzzy boarders of the shaded area are due to the stochastic nature of the single-cell data simulation - as discussed in the text, we have simulated a number of initially labelled cells high enough so that the simulation variance is negligible compared to the experimental uncertainty

were generated by simulating the system with a large sample of posterior parameter sets satisfying $p(\theta|\theta_{\text{MAP}}) \leq 0.05$ and then extracting the minimum and maximum resulting tumour sizes at each time point from the set of solutions.

#### T8.2 Induced knockdown of Tlx

It has previously been shown that a knock-out of Tlx has strong beneficial effects on animal survival in the tumour model discussed in this study [Zhu et al., 2014]. In the present work, a miRFlex system introduced via a viral vector was used to allow inducible knock-down of Tlx. Induction was then triggered by 10 consecutive injections of tamoxifen (TAM) and a CreERT2 excision system.

In order to model this experimental approach, we assume that each administration of TAM permanently turns a fraction $\epsilon$ of the current stem cell population into non-self-renewing progenitors of type $P$ irreversibly because of the reduction of active Tlx following the knock-down. The reason for a permanent instead of a transient switch lasting only for the time TAM is administered is that Cre drives an irreversible genetic change that remains after the end of TAM presence. However, because mRNA and protein have finite half-lives and the genetic switch is thus assumed to require some time until it takes effect, we assume that the conversion $S \longrightarrow P$ in the model does not happen instantaneously but with delay $\tau$. Otherwise, the equation system is preserved; cells behave as described in equations (4).

Main text figure 6F shows the model prediction resulting from this schedule for the maximum MAP parameter set $\theta^{\dagger}_{\mathrm{MAP}}$ and corresponding $\alpha = 0.05$ bounds when using conversion efficiencies of $\epsilon = 50\%$ and $\epsilon = 100\%$ respectively and a conversion delay of $\tau = 5\mathrm{d}$.
